# Retrotransposon addiction promotes centromere function via epigenetically activated small RNAs

**DOI:** 10.1101/2023.08.02.551486

**Authors:** Atsushi Shimada, Jonathan Cahn, Evan Ernst, Jason Lynn, Daniel Grimanelli, Ian Henderson, Tetsuji Kakutani, Robert A. Martienssen

## Abstract

Retrotransposons have invaded eukaryotic centromeres in cycles of repeat expansion and purging, but the function of centromeric retrotransposons, if any, has remained unclear. In *Arabidopsis*, centromeric *ATHILA* retrotransposons give rise to epigenetically activated short interfering RNAs (easiRNAs) in mutants in *DECREASE IN DNA METHYLATION1 (DDM1)*, which promote histone H3 lysine-9 di-methylation (H3K9me2). Here, we show that mutants which lose both DDM1 and RNA dependent RNA polymerase (RdRP) have pleiotropic developmental defects and mis-segregation of chromosome 5 during mitosis. Fertility defects are epigenetically inherited with the centromeric region of chromosome 5, and can be rescued by directing artificial small RNAs to a single family of *ATHILA5* retrotransposons specifically embedded within this centromeric region. easiRNAs and H3K9me2 promote pericentromeric condensation, chromosome cohesion and proper chromosome segregation in mitosis. Insertion of *ATHILA* silences transcription, while simultaneously making centromere function dependent on retrotransposon small RNAs, promoting the selfish survival and spread of centromeric retrotransposons. Parallels are made with the fission yeast *S. pombe*, where chromosome segregation depends on RNAi, and with humans, where chromosome segregation depends on both RNAi and HELLS^DDM1^.

## Main

Eukaryotic centromeres are usually composed of repetitive sequences with a unique chromatin composition that includes the centromeric histone H3 variant, CenH3^1^. CenH3 assembles the kinetochore, a large protein complex that attaches the chromosome to the spindle^1^. The positioning of CenH3 is thought to be epigenetically defined by surrounding pericentromeric heterochromatin – chromosomal material that remains condensed in interphase^2, 3^. Pericentromeric heterochromatin is also responsible for sister chromatid cohesion at mitosis, which ensures segregation of sister chromatids to each daughter cell during anaphase^1^. In most eukaryotes, these repetitive centromere sequences are composed of rapidly evolving tandem satellite repeats^1, 2^. In plants, many animals and fungi, satellite repeats are interspersed with specific classes of retrotransposons but the function, if any, of these retrotransposons has remained obscure^1^.

DNA methylation and RNA interference (RNAi) are important epigenetic pathways for both transcriptional and post-transcriptional gene silencing (TGS and PTGS). In the model plant *Arabidopsis thaliana*, DNA methylation is required to silence transposons, and can be triggered by RNAi through a pathway called RNA-dependent DNA methylation (RdDM). RdDM relies on 24-nt siRNAs produced by RNA POLYMERASE IV, RNA-DEPENDENT RNA POLYMERASE 2 (RDR2) and DICER-LIKE 3 (DCL3)^4, 5^. These 24-nt small RNAs bind to ARGONAUTE 4 (AGO4) and related proteins, which are thought to recruit DNA methyltransferases to RNA POLYMERASE V, along with other chromatin modifying enzymes^4, 5^. In organisms without DNA methylation, such as *Drosophila*, *Caenorhabditis elegans* and the fission yeast *Schizosaccharomyces pombe*, RNAi guides histone modifications, notably dimethylation of histone H3 lysine-9^6–8^, which plays a major role in centromere cohesion. For this reason *S. pombe* mutants in RNAi have strong defects in chromosome segregation^9, 10^. In Arabidopsis, such mitotic defects are very mild, or not apparent when components of the canonical RdDM pathway are mutated, despite the complete loss of 24-nt small RNA^11^.

DNA methylation can be maintained in the absence of RdDM by the DECREASE IN DNA METHYLATION1 (DDM1) SWI2/SNF2 chromatin remodeler, but mutants retain fertility and normal chromosome segregation despite substantial demethylation of centromeric satellite repeats^12^. While RdDM-mediated DNA methylation is required for TGS, Arabidopsis possesses another RNAi pathway for PTGS, which generates 21-nt or 22-nt siRNAs via *RDR6*-*DCL2/DCL4-AGO1/AGO7* and silences euchromatic genes, transgenes and viral RNAs^13^. We previously identified a new class of epigenetically activated 21-nt siRNAs (easiRNAs) derived from transposable elements in *ddm1* mutants^14^, which have elevated transcription of transposons^15^. Similar small RNAs are found in *ddm1-like* double mutants in maize, although mutant embryos fail to germinate in this species^16^. We rationalized small RNAs might compensate for the loss of DNA methylation in *ddm1* mutants, and set out to determine the developmental and chromosomal consequences of removing RNAi in the absence of DNA methylation.

### Epigenetic defects at Arabidopsis centromeres in RNAi and DNA methylation mutants

The biosynthesis of 21-22nt easiRNAs is dependent on RDR6^17, 18^, which is partially redundant with RDR1^19^, while RDR2 contributes to RNA-dependent DNA methylation via 24nt siRNAs^20, 21^. We have previously shown that *ddm1* mutants bearing mutations of all three RNA-dependent RNA polymerase genes *rdr1 rdr2 rdr6 ddm1* (hereafter *rdr1;2;6 ddm1)*, have severe developmental defects, unlike *ddm1*, *rdr1;2 ddm1* or *rdr1;2;6* alone^17, 18^. *rdr1;2;6 ddm1* mutants exhibit pleiotropic developmental defects such as infertility, short stature, slow growth, curly leaves and flowers with additional stamens and missing organs (Fig. 1a and Extended Data Fig. 1a). By contrast, no conspicuous phenotype is observed in *rdr1;2 or rdr1;2;6* mutants, while in *rdr1;2; ddm1* mutants only vegetative phenotypes, such as curly leaves and short stature, are observed (Fig. 1a and Extended Data Fig. 1a). Thus *RDR6* activity is essential for fertility and floral organ differentiation in the absence of DNA methylation^18, 22^. Importantly, backcrosses to *rdr1;2;6* triple mutants demonstrated that these phenotypes were inherited epigenetically when *DDM1* function was restored in the absence of RDRs (Fig. 1b).

**Figure 1.**
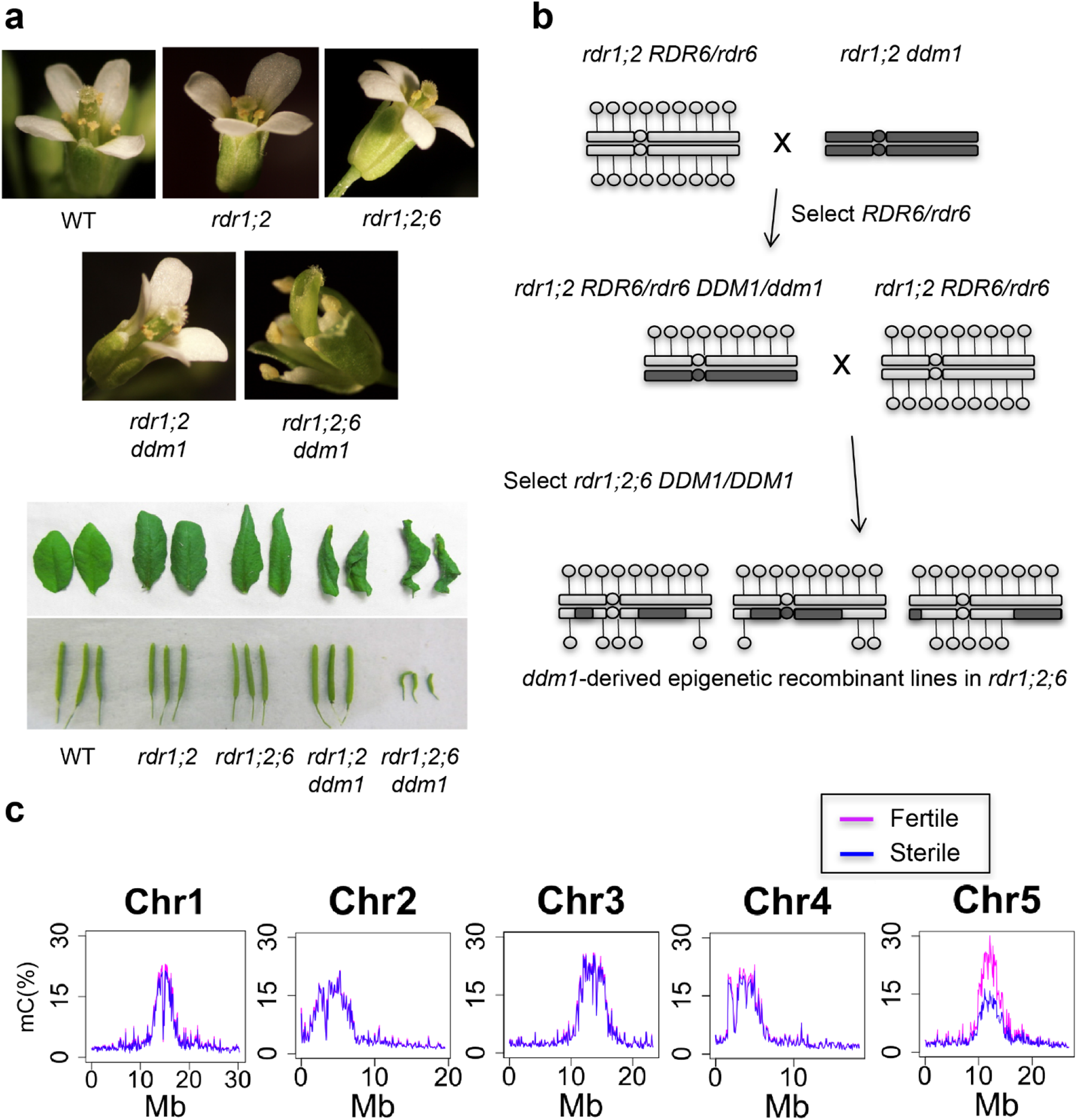
Fertility and floral defects of *rdr1;2;6 ddm1* map to hypomethylated centromere 5. **a**, Developmental defects of double, triple and quadruple mutants in RNA dependent RNA polymerase (*rdr1*, *rdr2*, *rdr6*) and DNA methylation (*ddm1*) in floral organ identity, leaf shape, and fertility (silique length). *rdr1 rdr2* and *rdr6* combinations are abbreviated as *rdr1;2;6*. **b**, Crossing scheme for constructing ddm1-derived epigenetic recombinant lines in an *rdr1 rdr2 rdr6* background. Hypomethylated chromosomal regions derived from *ddm1* mutants are inherited epigenetically in DDM1/DDM1 progeny, and are indicated in dark grey. **c**, Whole genome bisulfite sequencing of pooled fertile (pink) and sterile (blue) epigenetic recombinant lines (from **b**) indicates reduced methylation in the pericentromeric regions of chromosome 5.

Because mutants defective in DNA methylation have been shown to suffer from developmental phenotypes due to mis-expression of individual genes^23–25^, we hypothesized that there might be a causative locus which is silenced by *RDR6*-dependent 21-nt siRNAs in *ddm1* mutants. To identify this locus, we performed genetic mapping by generating *ddm1*-derived epigenetic recombinant lines in an *rdr1;2;6* mutant background. Because the loss of DNA methylation in *ddm1* is epigenetically inherited, especially in an *rdr2* mutant background^21^, *ddm1*-derived chromosomes remain hypomethylated even after backcrossing to WT. This allowed us to identify which chromosomal region(s) was responsible for the phenotype. Epigenetic recombinant lines in an *rdr1;2;6* background were generated in the crossing scheme shown in Fig. 1b. Plants were classified into 4 groups depending on their phenotypes, namely (1) WT-like, (2) curly-leaf, (3) sterile and (4) both sterile and curly leaf (Extended Data Fig. 1b-g). The sterile phenotype was always associated with the floral defect (Extended Data Fig. 1b-g), suggesting that these defects may arise from the same dominant mutation. We performed whole genome bisulfite sequencing analysis to compare genome-wide DNA methylation levels between sterile and fertile plants from these backcrosses. This analysis demonstrated that the sterile and floral phenotypes were linked to the hypomethylated centromeric region of chromosome 5, derived from *ddm1* (Fig. 1c). We performed fine mapping using McrBC, a restriction enzyme digesting only methylated DNA, and amplification by PCR, to determine whether a given chromosomal region was *ddm1*-derived or WT-derived^26^. In this way, the causative locus was mapped to the interval between AT5G28190 and AT5G36125 (Extended Data Fig. 2). Although we examined more than 200 individuals, we could not narrow down this causative interval further because of the low frequency of meiotic crossovers in centromeric regions^27^.

**Figure 2.**
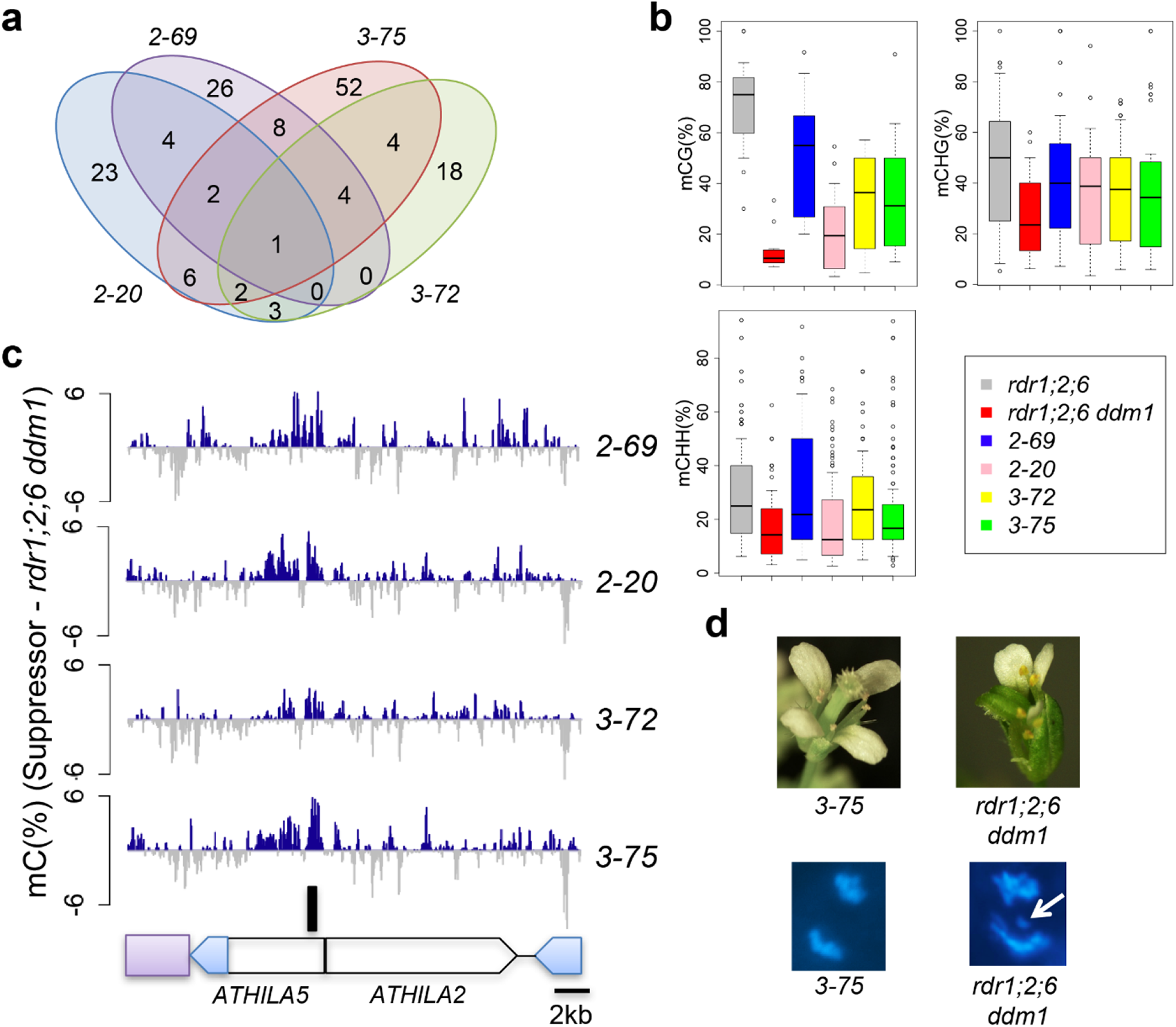
Epiallelic suppressors gain DNA methylation at the *ATHILA5* retrotransposon in centromere 5. **a**, Venn diagram of shared, hypermethylated differentially methylated regions (DMR) in 4 independent *rdr1;2;6 ddm1* suppressors (*2-69*, *2-20*, *3-72*, *3-75*), on chromosome 5. **b**, Boxplot analyses of DNA methylation levels in uniquely shared 1kb hyper-methylated DMR in each genotype. **c**, Uniquely shared DMR (black bar) corresponds to *Cen5-ATHILA5* (blue LTRs) which is embedded in cen180 satellite repeats (purple box) and interrupted by *ATHILA2*. Genome Browser tracks display DNA methylation gains (blue) and losses (grey) in the 26kb region relative to *rdr1;2;6 ddm1*. **d**, Floral and chromosomal phenotypes of *rdr1;2;6 ddm1* mutants are rescued by epiallelic suppressor *3-75*. Mitotic chromosomes in root tip anaphase cells were stained with DAPI. A mis-segregating chromosome is indicated (white arrow).

### An *ATHILA5* retrotransposon promotes centromere function on chromosome 5

Taking an alternative approach, we next performed EMS mutagenesis to obtain suppressors which rescue the fertility defect in *rdr1;2;6 ddm1* mutants. Four suppressors were isolated which also rescued the short stature and floral developmental defects (Extended Data Fig. 3a). Whole genome sequencing of pooled sterile and fertile segregants revealed that the suppressors were linked to single nucleotide polymorphisms (SNPs) on centromere 5 (Extended Data Fig. 3b), but curiously, there were no commonly mutated genes among the four suppressors and most of the introduced SNPs were in transposable elements (Extended Data Fig. 3c, d). Along with nucleotide substitutions, EMS mutagenesis is also capable of inducing changes in cytosine methylation, resulting in epialleles^28–30^. EMS-induced epialleles of *superman,* for example, gain DNA methylation in the promoter region and behave like *superman* mutants without any change in the DNA sequence. This led us to consider the possibility that suppression might be caused by epigenetic modification rather than nucleotide substitution. We performed whole genome bisulfite sequencing of pooled fertile and sterile segregants and DMR (Differentially Methylated Regions) analysis using the TAIR10 assembly of the Arabidopsis Col-0 genome revealed a single hyper-methylated locus in the interval on centromere 5 common to all four suppressors (Fig. 2a, b). This locus corresponds to the 5’ region of the *ATHILA5* retrotransposon *Cen5-ATHILA5* (Fig. 2c).

**Figure 3.**
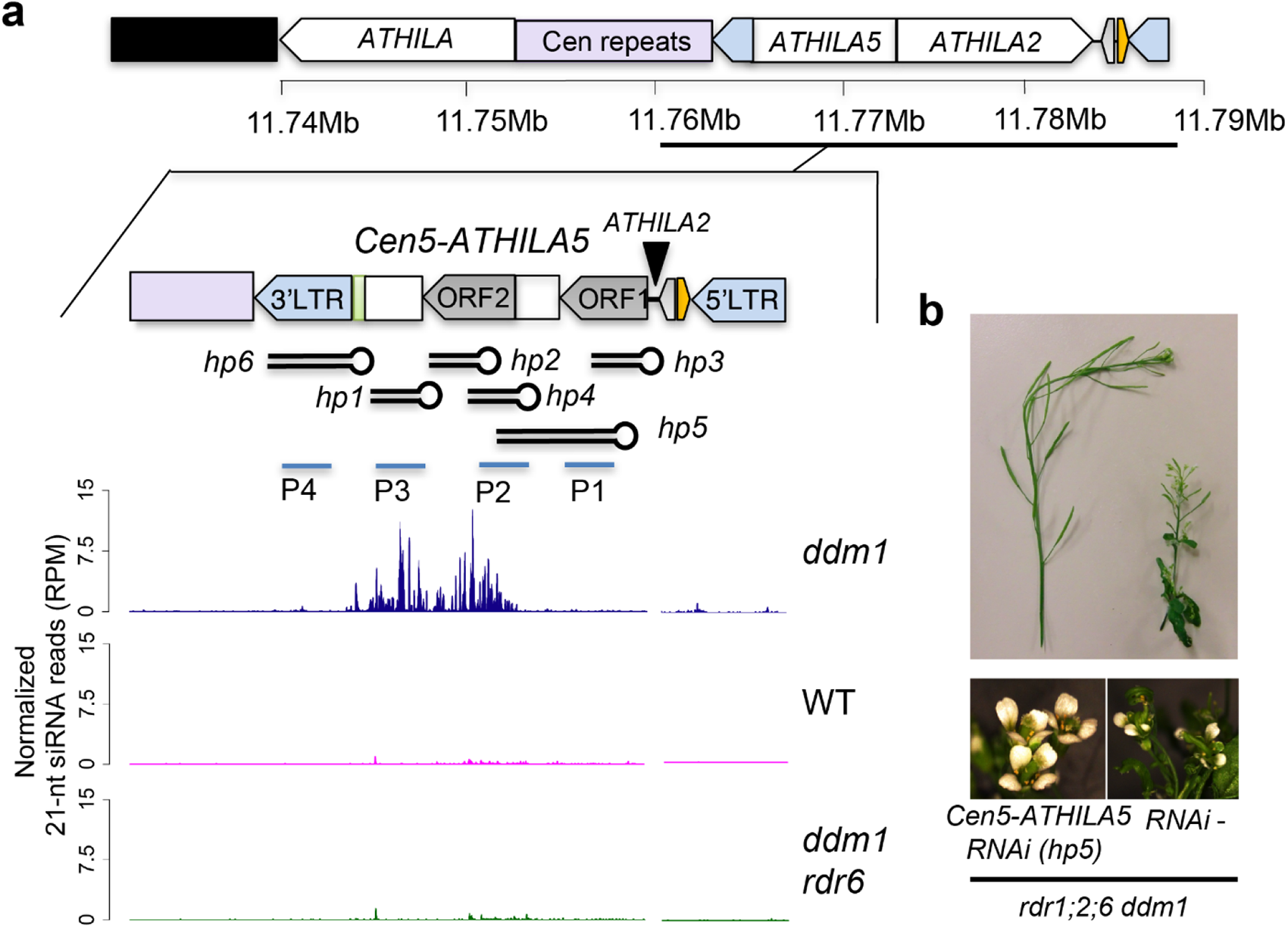
*Cen5-ATHILA5* hairpin small RNA rescue *rdr1;2;6 ddm1* phenotypes. **a**, *Cen5-ATHILA5* is embedded in sequenced (purple) and unsequenced (black) centromeric repeats on chromosome 5 (TAIR10 genome assembly). It encodes two ORFs (gray arrows), short regions of homology to mitochondrial DNA (light green) and a tRNA gene (orange). Synthetic hairpins hp1 through hp6 and probes used for northern analysis (Extended Data Fig. 5) are shown. Genome browser tracks display 21-nucleotide siRNA levels in indicated genotypes. **b**, RNAi hairpin hp5 strongly suppresses floral and fertility defects in *rdr1;2;6 ddm1*.

The vegetative and infertility phenotypes of *rdr1;2;6 ddm1* mutants strongly resemble the phenotypes of plants expressing centromeric histone cenH3 “tailswap” GFP fusions, in which centromere function is impaired^31^. Therefore, we examined root tip anaphase cells in each of the genotypes for lagging chromosomes, an indication of impaired centromere function^31^. Remarkably, there was a strong chromosome lagging phenotype in *rdr1;2;6 ddm1* mutants, but not in the other genotypes (Fig. 2d; Table 1). This phenotype was ameliorated to some extent in the epigenetic suppressors (Table 1), although visible phenotypes returned in the next generation, consistent with the instability of these epialleles in a *ddm1* background. This data strongly suggested that centromere function was disrupted in *rdr1;2;6 ddm1* mutants, and was epigenetically inherited in the absence of RNAi (Fig. 1b).

**Table 1.** Mitotic chromosome mis-segregation in *rdr1;2;6 ddm1*. Chromosome mis-segregation was observed in root tip mitotic cells (n=100) and the mis-segregation rate was calculated in the indicated strains. *tailswap cenh3*, a mutant defective in kinetochore function^31^, was used as a positive control.

|  | Mis-segregation |
| --- | --- |
| WT | 0 % |
| *rdr1;2* | 0 % |
| *rdr1;2;6* | 0 % |
| *rdr1;2 ddm1* | 0 % |
| *rdr1;2;6 ddm1* | 31 % |
| *rdr1;2;6 ddm1 hp5 RNAi* | 3 % |
| *tailswap cenh3* | 19 % |
| episuppressor 3-75 | 18 % |
| *rdr1;2 ddm1 kyp* | 13 % |
| *rdr1;2 kyp* | 0 % |

### Retrotransposon small RNAs rescue defects in chromosome segregation

*Cen5-ATHILA5* encodes AT5G31927, which comprises two open reading frames, the GAG gene (ORF1) and an *ATHILA* superfamily gene (ORF2), but ORF1 is interrupted by the integration of another retrotransposon, *ATHILA2,* potentially rendering it incompetent for further transposition (Fig. 3a). The expression level of *Cen5-ATHILA5* is higher in *rdr1;2;6 ddm1* than in *rdr1;2 ddm1* or *rdr1;2;6,* and was silenced in the suppressor mutants (Extended Data Fig. 4a). We first hypothesized that proteins coded by *Cen5-ATHILA5* might be responsible for the mutant phenotype, but overexpression of the entire *Cen5-ATHILA5* element or ORF AT5G31927 did not cause any phenotype in *rdr1;2;6* mutant backgrounds (Extended Data Fig. 4b, c). Instead we considered the possibility that the loss of easiRNAs might be responsible as *Cen5-ATHILA5* 21-nt easiRNAs accumulate in *ddm1*, but not in *ddm1 rdr6* (Fig. 3a)^18^. Simple overexpression of *Cen5-ATHILA5* would not be expected to restore easiRNAs in the absence of RNA dependent RNA polymerase, so instead we introduced *Cen5-ATHILA5* hairpins into the *rdr1;2;6 ddm1* mutant as a source of double stranded easiRNAs independent of RdRP (Fig. 3a). Hairpins corresponding to *ATHILA2* easiRNAs were also introduced as controls. These hairpins all generate 21-nt and 24-nt small RNAs (Extended Data Fig. 5a).

**Figure 4.**
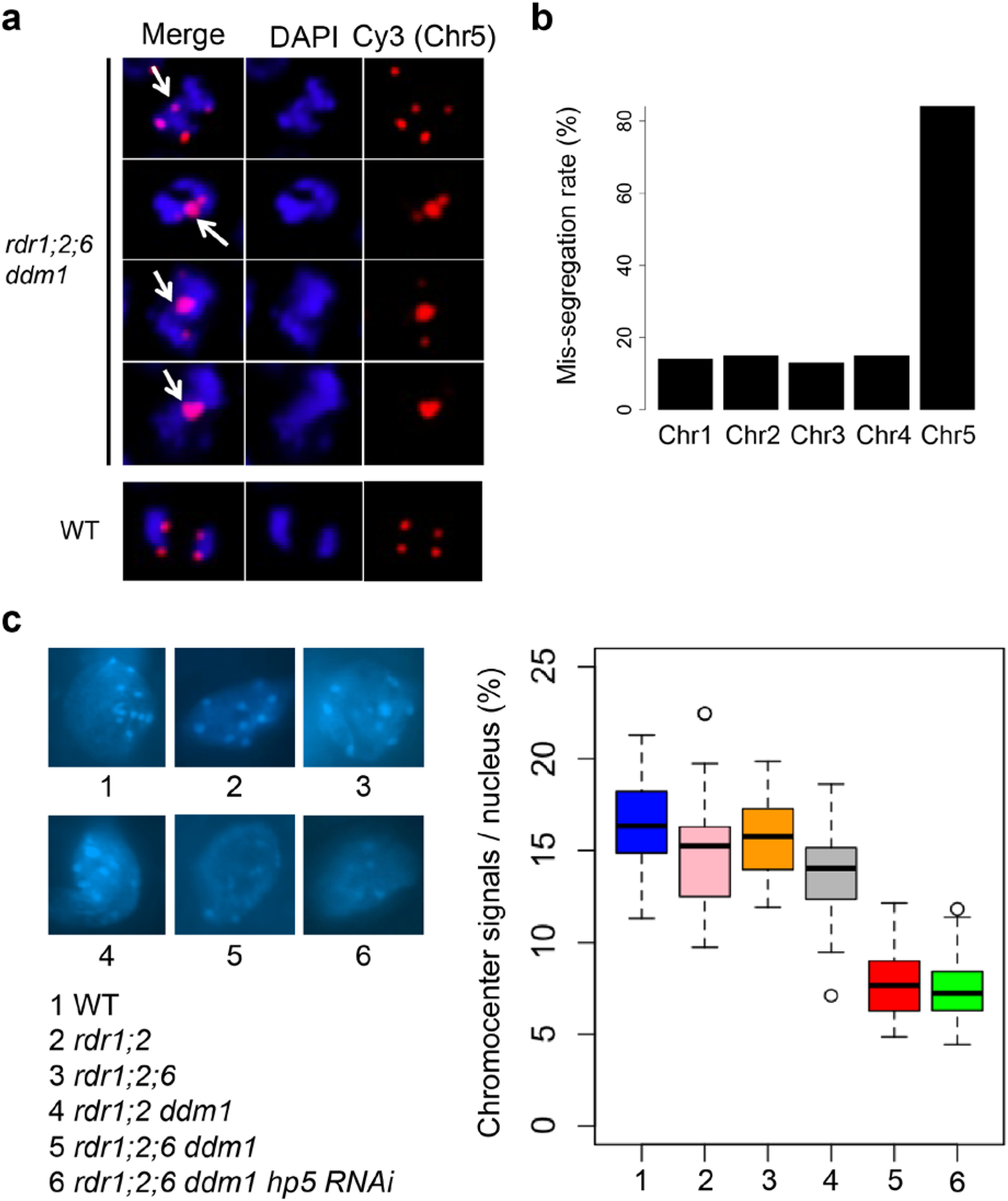
Chromosome mis-segregation in *rdr1;2;6 ddm1*. **a**, DNA FISH of root tip anaphase cells with Cy3-labeled DNA probes from chromosome 5 (red). Nuclei were counterstained with DAPI. Mis-segregating chromosomes are indicated with white arrows. **b**, Mis-segregation rate of chromosomes 1-5 in *rdr1;2;6 ddm1* anaphase cells (n=100 abnormal cells) determined by FISH. **c**, Chromocenters were stained with DAPI, and quantified signals (n=30) illustrated by boxplots (right). 1, WT; 2, *rdr1;2*, 3 *rdr1;2;6*, 4, *rdr1;2;6 ddm1*, 5 *rdr1;2;6 ddm1 hp5 RNAi*.

**Figure 5.**
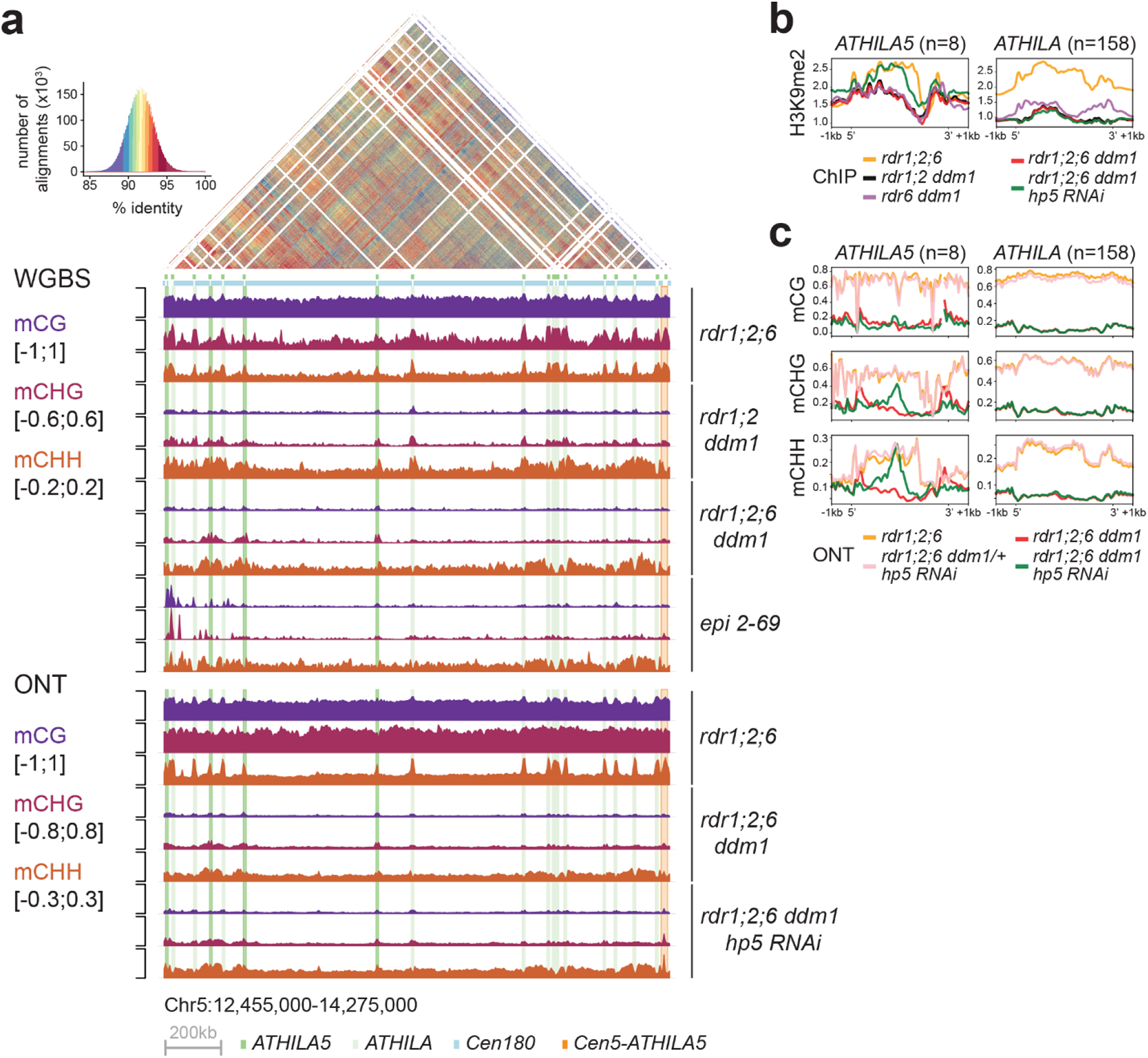
easiRNAs restore H3K9me2 and non-CG DNA methylation to *ATHILA5* elements of chromosome 5. **a**, Browser shot of DNA methylation in the different mutants, spanning the centromere of Chromosome 5 (Col-cen genome assembly). The dotplot reveals high identity between the cen180 repeats (light blue bar), with interspersed *ATHILA* elements (light green columns) and *ATHILA5* elements (dark green columns). The *CEN5-ATHILA5* used to design the RNAi hairpin is also shown (orange column, far right). DNA methylation was assessed by whole-genome bisulfite sequencing (WGBS), as well as by base calling from long-reads from Oxford Nanopore (ONT) using modbase2. mC calls are averaged in 5kb windows (y axis scale indicated for each context). **b**-**c**, Metaplots of (**b**) H3K9me2 and (**c**) DNA methylation (from ONT) over *ATHILA5* (not including hp5 containing *CEN5-ATHILA5*) and all other *ATHILA* in the genome. Levels of H3K9me2 and non-CG methylation in the *rdr1;2;6 ddm1* mutant are recovered specifically at *ATHILA5* elements when the hp5 is present.

Remarkably, hairpin-derived *Cen5*-*ATHILA5* small RNAs corresponding to both *ORF1* and *ORF2* (*hp5*) rescued infertility and many of the pleiotropic developmental defects (Fig. 3b and Extended Data Fig. 5b), while slightly milder suppression was observed for hairpins (*hp2,4*) targeting *ORF2* alone (Extended Data Fig. 5c). All 3 rescuing hairpins overlap with the easiRNA-accumulating region. This suppression was not observed when hairpins matching *ATHILA2* (including *hp1*, *hp7* and *hp8*) were introduced (Extended Data Fig. 5c-f). Most importantly, the high frequency of mitotic chromosome mis-segregation in the *rdr1;2;6 ddm1* mutant was also greatly reduced in the *Cen5-ATHILA5* hairpin *hp5* suppressor (Table 1). Because *Cen5-ATHILA5* is embedded within the 180bp centromeric satellite repeats of chromosome 5 (Fig. 3a), the mis-segregating chromosomes observed in the *rdr1;2;6 ddm1* mutant should correspond to chromosome 5. To test this hypothesis, we performed DNA FISH using mitotic cells from root tips in the *rdr1;2;6 ddm1* mutant. The proportion of mis-segregating chromosome 5 was calculated by co-localization of Cy3 probe signals with the observed chromosomal mis-segregation. 84% of mis-segregating chromosomes correspond to chromosome 5, with lower proportions for the other chromosomes (Fig. 4a, b). Thus artificial siRNAs derived from the *Cen5-ATHILA5* retrotransposon are sufficient to restore accurate chromosome segregation.

### Retrotransposon small RNAs promote DNA methylation and histone H3K9 di-methylation

Recently, the centromeric sequences of Col-0 have been assembled with single molecule long-read sequencing technology^32^. Unexpectedly, multiple copies of intact *ATHILA5* and other *ATHILA* retrotransposons were specifically found embedded into the cenH3-containing centromeric repeats of centromere 5. Single molecule long read sequencing and assembly of other Arabidopsis accessions have since revealed that waves of *ATHILA5* retrotransposons have recently and specifically disrupted centromeres 4 and 5 in several sympatric accessions of Arabidopsis from Europe^33^. Invasion seems to have disrupted homogenization of satellite repeats, suggesting these insertions may interfere with recombination mechanisms such as break induced replication and repair^33^.

Mapping of our whole genome bisulfite sequencing data revealed that satellite repeats lost CG and CHG methylation in *ddm1* mutant combinations, but retained CHH methylation as expected. However, the *ATHILA* elements in centromere 5 retained some CHG and especially CHH methylation in *rdr1;2 ddm1,* but substantially less in *rdr1;2;6 ddm1.* Furthermore, WGBS data from a heritable epigenetic suppressor (2-69, Fig. 2a, b) also revealed ectopic DNA methylation at *ATHILA* elements, but in all 3 sequence contexts (Fig. 5a). In order to detect methylation at cytosine residues unambiguously in highly repetitive regions, we performed single molecule very long read genome sequencing using Oxford nanopore technology (ONT), and the methyl cytosine base-calling protocol (see Methods). We compared methylation patterns in *rdr1;2;6 ddm1*, and in *rdr1;2;6 ddm1/+* siblings, with and without the *Cen5-ATHILA5* (*hp5)* hairpin suppressor (Fig. 5a). On metaplots of *ATHILA5* elements, but not other *ATHILA* elements, DNA methylation was specifically restored precisely in the region defined by the hairpin. DNA methylation was only restored in the CHG and CHH contexts, and not in the CG context, consistent with being induced by RNAi (Fig. 5c).

CHG and CHH DNA methylation depend on histone lysine-9 di-methylation (H3K9me2) via the chromodomain DNA methyltransferases CHROMOMETHYLTRANSFERASE2 (CMT2) and CMT3. In *Schizosaccharomyces pombe*, RNAi mutants lose H3K9me2 and suffer from severe chromosome mis-segregation due to loss of sister chromatid cohesion^9, 10^. In Arabidopsis *ddm1* mutants, *RDR6*-dependent easiRNAs derived from pericentromeric transposons also induce H3K9me2^34^, and we postulated that they might have a role in centromeric organization. We observed that chromocenters in *rdr1;2;6 ddm1* were greatly diminished when compared with those in *rdr1;2 ddm1*, or *rdr1;2;6* mutants (Fig. 4c), suggesting that *RDR6* activity, specifically in the absence of DNA methylation, is required for pericentromeric heterochromatin condensation. We then investigated the effect of *rdr6* on histone modification by comparing *rdr1;2;6* and *rdr1;2;6 ddm1* mutants with and without the addition of *hp5*. We performed chromatin immunoprecipitation sequencing (ChIP-seq)^35^ and found that H3K9me2 is highly enriched in multiple families of *ATHILA* elements in WT, but reduced in *rdr1;2 ddm1* and *rdr1;2;6 ddm1* (Fig. 5b), a result which was confirmed by immunofluorescence (Extended Data Fig. 7). However, this decrease in H3K9me2 was almost fully restored by the *Cen5-ATHILA5* hairpin suppressor along with chromosome segregation (Fig. 5b). Thus, easiRNAs ensure pericentromeric H3K9me2 at *ATHILA5* elements in centromere 5. Severely diminished chromocenters, sterile and developmental phenotypes are also observed when Arabidopsis loses both histone H3K9 and DNA methylation^36^. We further tested this idea by making mutant combinations with *KRYPTONITE*, one of several H3K9 methyltransferases in Arabidopsis. We found that vegetative phenotypes of *rdr1;2 ddm1 kyp* mutants resembled *rdr1;2;6 ddm1* mutants, including defects in chromosome segregation, although floral defects were less severe (Extended Data Fig. 8).

Pericentromeric heterochromatin near the kinetochore includes the inner centromere, which connects sister kinetochores before anaphase through chromosome cohesion. In mammals and yeast, the inner centromere functions as a scaffold to recruit factors important for chromosome segregation such as Aurora kinase, shugoshin, cohesin and condensin^37^. Although the Arabidopsis inner centromere has not been well characterized, cohesin and condensin are enriched in the pericentromere^38, 39^, and mutation of these factors affects pericentromeric architecture and chromosome mis-segregation^39, 40^. Further, histone residues H3S10 and H3T3 are highly phosphorylated specifically at the pericentromeric region during mitosis^41^, and the activity of Aurora kinase is essential for chromosome segregation^42^. We examined H3T3 phosphorylation by antibody staining, and could clearly detect phosphorylation at chromocenters, which were smaller in *rdr1;2;6 ddm1* than *rdr1;2 ddm1* as expected (Fig. 6a). Next, we used DNA FISH of chromosome 5 to examine cohesion in the mutants. By counting the number of dots, we could assess whether cohesion was normal at mitosis (2 dots) or reduced (3 or 4 dots). We found that cohesion was dramatically lost in *rdr1;2;6 ddm1* mutants, but fully restored by the *Cen5-ATHILA5* (hp5) hairpin (Fig. 6b).

**Figure 6.**
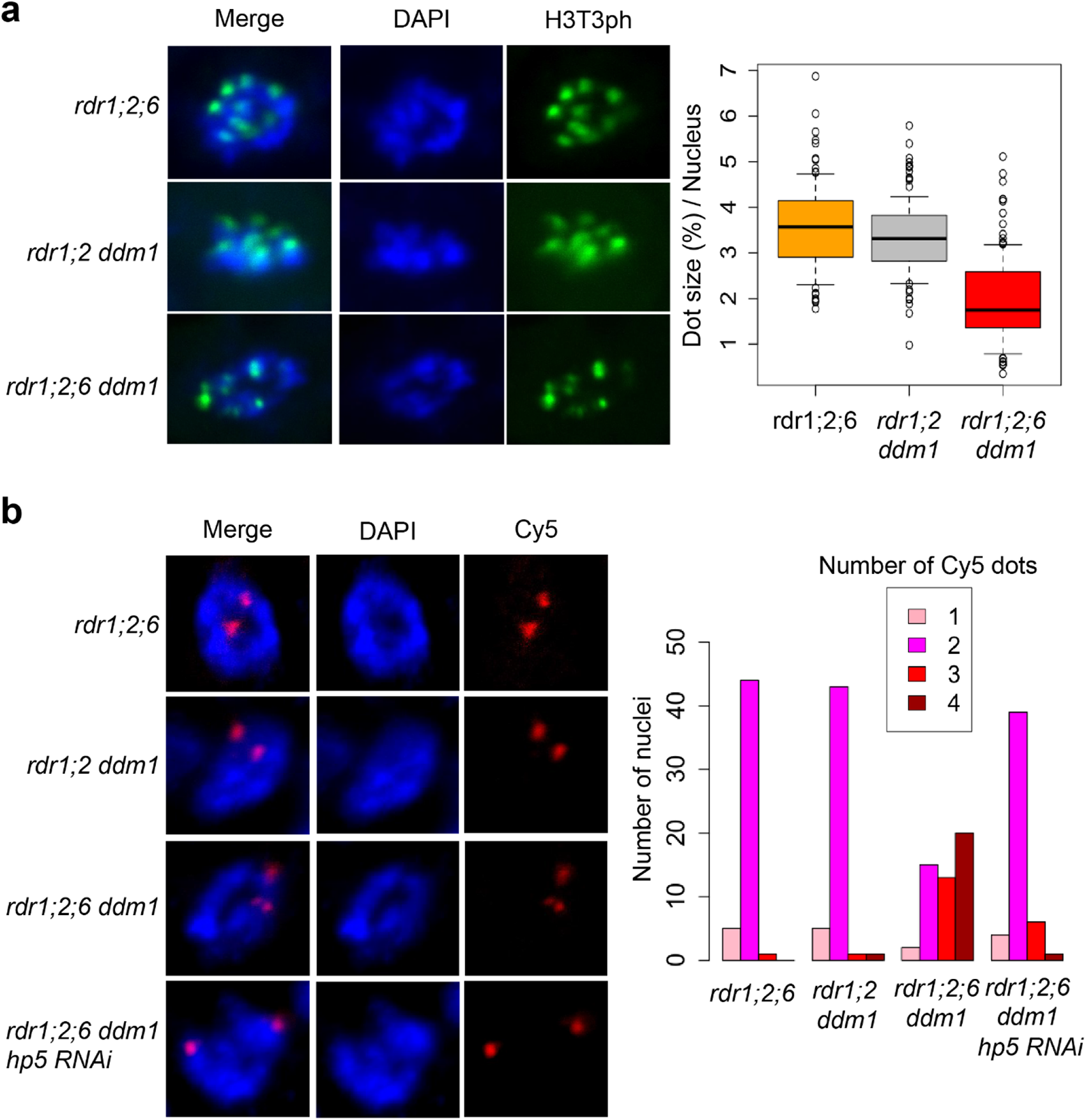
Defective sister chromatid cohesion in *rdr1;2;6 ddm1*. **a**, Immunofluorescence for H3T3ph in root tip cells. Left panels exhibit mitotic prophase cells showing H3T3ph signals. Cells were counterstained with DAPI. In prophase cells, condensed DAPI dots are dispersed in the nucleus. H3T3ph dot sizes were calculated based on the nucleus size (Right panel). 100 dots from 20 nuclei were analyzed. **b**, DNA FISH in mitotic prophase cells with Cy5-labeled probes designed near the pericentromeric region of chromosome 5 (Left Panels). The right panel shows the number of Cy5 dots (1-4) in each nucleus. 50 nuclei were analyzed for each mutant.

### RNAi and histone modification guide chromosome cohesion in Arabidopsis

We demonstrate that *RDR6*-dependent 21-nt easiRNAs compensate for loss of DNA methylation by promoting pericentromeric chromatin condensation and proper mitotic chromosome segregation (Fig. 7). We did not examine meiotic chromosome segregation due to the difficulty of identifying meiotic cells in quadruple mutants, and it is likely that developmental defects may account for their near-complete infertility (Extended Data Fig. 9). We observed that *RDR6*-dependent 21-nt easiRNAs facilitate histone H3K9 methylation in the absence of DDM1, and are required for chromosome segregation and normal development. Importantly, the phenotypic defect in *rdr1;2;6 ddm1* was rescued by restoring small RNAs and histone H3K9 methylation via hairpin precursors that match *Cen5-ATHILA5,* a Ty3/gypsy class retrotransposon family embedded specifically within *Cen5* centromeric repeats. Similar hairpin precursors induce H3K9me2 and non-CG DNA methylation in Arabidopsis^43, 44^. However, we did not observe any difference in H3K9me2 levels between *rdr6 ddm1, rdr1;2 ddm1* and *rdr1;2;6 ddm1* mutants (Fig. 5b), despite having differing levels of chromosome segregation (Fig 6). As *rdr1;2 kyp ddm1* mutants resemble *rdr1;2;6 ddm1* mutants in this respect, we speculate that an additional histone modification is likely guided by 24nt siRNAs, mediated by *RDR2*. Both modifications are presumably required for cohesion.

**Figure 7.**
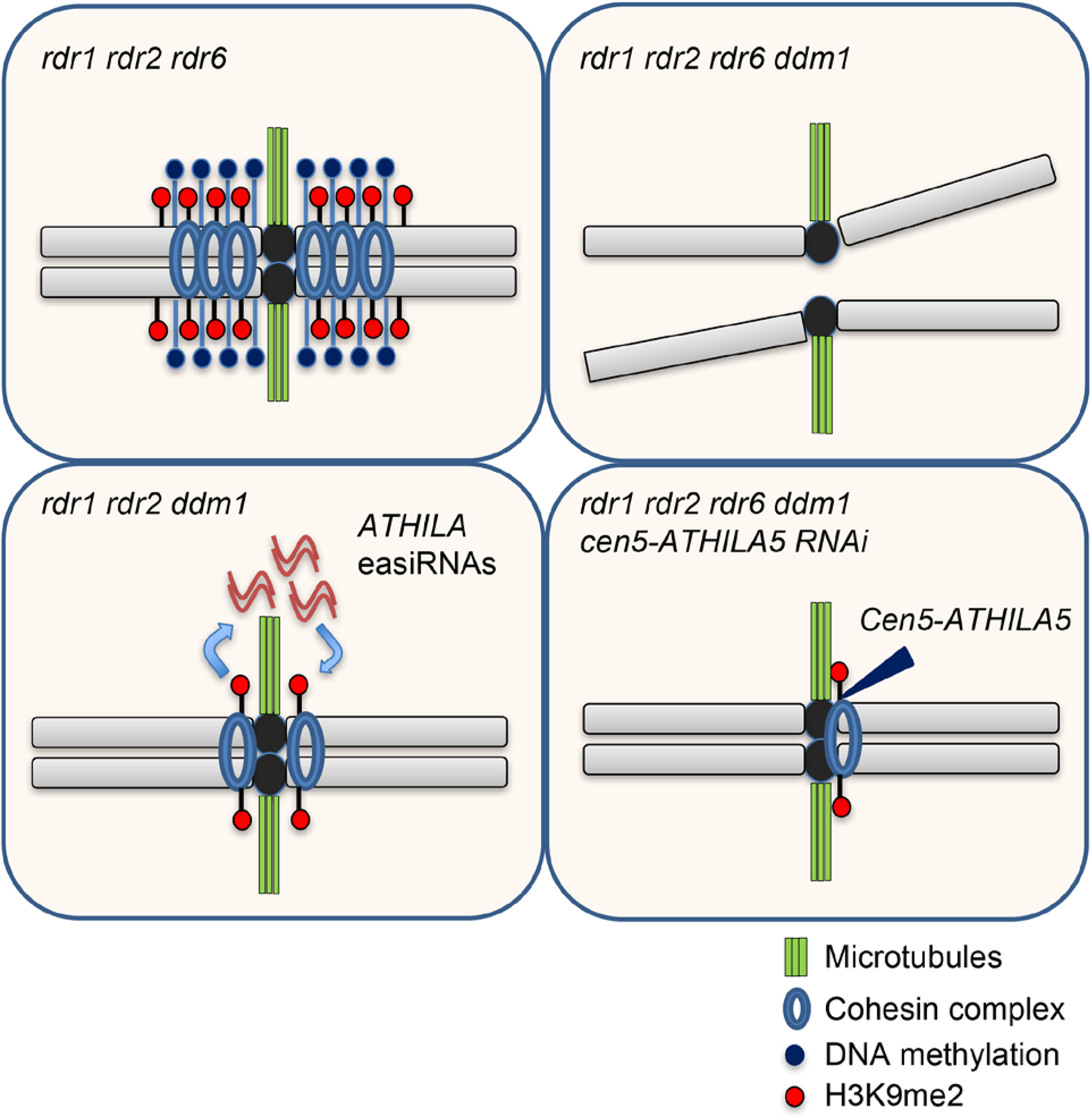
A model for the regulation of pericentromeric sister chromatid cohesion by DNA methylation and small RNAs. The Arabidopsis pericentromere is maintained by DNA methylation and H3K9 methylation, and is essential for sister chromatid cohesion. When DNA methylation is lost, plants produce RDR6-dependent easiRNAs from *ATHILA* family retrotransposons, enriching H3K9 methylation at pericentromeric *ATHILAs*. Additional loss of easiRNAs causes impaired sister chromatid cohesion and severe mis-segregation of chromosome 5. The sterility, and the sister chromatid cohesion defect of chromosome 5 can be rescued by artificial small RNAs targeting retrotransposon *ATHILA5*, which re-establishes H3K9-methylated heterochromatin.

In the fission yeast *S. pombe*, which lacks DNA methylation, RNAi promotes sister chromatid cohesion by recruiting cohesin to pericentromeric heterochromatin and allowing proper chromosome segregation^9, 10^. In mouse, *Dicer* mutant ES cells also have strong centromeric segregation defects, and these can be rescued by mutations in conserved transcription factors that also rescue *dicer* mutants in fission yeast^45^. In humans, patients with ICF syndrome (immunodeficiency, centromere function and facial abnormalities) have mutations in HELLS, the DDM1 ortholog, or in other genes required for DNA methylation, and HEK293 cells mutant for these genes have defects in chromosome segregation and DNA methylation^46^. This suggests that mammalian cells require both RNAi and DNA methylation for centromere function. In *Arabidopsis,* we show that chromosome segregation can be maintained by either RNAi or DNA methylation, so that only mutants that lose both have segregation defects. In each species, including mammals and plants^11, 45^, these effects are likely mediated by centromeric transcription which is silenced by histone H3K9 methylation, promoting cohesion. Humans lack RNA dependent RNA polymerase, which amplifies siRNAs in yeast and Arabidopsis, and loss of HELLS^DDM1^ alone leads to immune and centromere defects. Perhaps siRNAs targeted to centromeric repeats offer a potential therapeutic avenue for ICF syndrome^47^.

### Retrotransposon invasion and centromere function

While segregation of all 5 chromosomes was defective in *rdr1;2;6 ddm1,* mis-segregation of chromosome 5 had the largest phenotypic contribution, and co-segregated epigenetically with the loss of DNA methylation in this interval, strongly supporting the idea that centromere function is an epigenetic property^48^. We note that trisomics of chromosome 5, among all the Arabidopsis trisomics, exhibit the most severe defects in fertility^49^, which might explain why fertility defects mapped to this centromere in particular. The Arabidopsis inner centromere is comprised of tens of thousands of 180bp repeats, but Col-0 chromosome 5 stands out in having been recently invaded by *ATHILA* retrotransposons, notably by *ATHILA5*^32^. It has previously been reported that a subset of centromeric satellite repeats are transcribed but post-transcriptionally silenced by DCL1, which triggers easiRNAs^11, 17^. Sequence comparison indicates that these repeats bind cenH3^50^. Another subset of repeats is transcriptionally silenced by DDM1, which prevents transcription from embedded *ATHILA* retrotransposons and their derivatives^11^. Both classes are associated with DNA methylation and H3K9me2^32^. Thus, in addition to centromere disruption, insertion of centromeric *ATHILA* retrotransposons silences transcription from centromeric repeats by a combination of DNA methylation, RNAi, and H3K9me2. The loss of centromere function at centromeres that have been recently disrupted and silenced, suggests that retrotransposon invasion simultaneously makes centromeres dependent on these very same transposons. Thus, centromeres become “addicted” to invading retrotransposons via RNAi and silencing^51^, an apparently successful strategy for retrotransposon survival in Arabidopsis^33^. Similar strategies may have been deployed in maize^52^ and in a close fission yeast relative^53^ whose centromeres have also been recently invaded.

## Methods

### Plant strains, preparation of DNA and RNA, primers

*ddm1-1* mutants with mutations of RNA dependent RNA polymerase genes (*rdr1* (SALK_112300) *rdr2* (SALK_059661) *rdr6-11*) were generated in the previous study^22^. *tailswap cenh3* is a kind gift from Dr. Simon W. L. Chan. The *kyp-4* (SALK_044606) mutant was used.

DNA was extracted from leaves of 4 week-old plants by Nucleon Phytopure (GE Healthcare) and total RNA was extracted from 3 week-old plants by RNeasy (QIAGEN) or Direct-zol (ZYMO RESEARCH). Primers used in this study are listed in Supplementary Table 1.

### Construction of “epi-recombinant lines”

*rdr1 rdr2 ddm1* was crossed to *rdr1 rdr2 RDR6/rdr6* to obtain *rdr1 rdr2 DDM1/ddm1 RDR6/rdr6* plants in F1. The F1 *rdr1 rdr2 DDM1/ddm1 RDR6/rdr6* plants were crossed to *rdr1 rdr2 RDR6/rdr6*. *rdr1 rdr2 rdr6/rdr6 DDM1/DDM1* were selected in F2 and DNA were extracted from the rosette leaves of 4 week-old plants individually (10 fertile and 10 sterile plants), followed by whole genome bisulfite sequencing as described previously^18^. DNA methylation levels in all three cytocine contexts (CG, CHG and CHH) in 100kb fixed windows were calculated for each sample, and the average DNA methylation levels for the fertile and sterile groups were compared.

### Hairpin small RNA complementation

The 35S promoter and nos terminator were cloned into pPZP2H^54^ to make an expression vector (p35S-pPZP2H) at KpnI-ApaI and XbaI-SacI site, respectively. A partial *Cen5-ATHILA5* element and its inverted form separated with *GUS* spacer were amplified by PCR (T8H11 BAC DNA and *Saccharomyces cerevisiae* genomic DNA were used as templates to amplify *Cen5-ATHILA5* and *GUS* fragments) and cloned into p35S-pPZP2H, resulting in inverted repeats of *Cen5-ATHILA5* in the expression vector. After transformation using *Agrobacterium tumefacien*s, *DDM1/ddm1* T1 transformants were selected with hygromycin resistance, and the T2 plants were grown without hygromycin selection for each hairpin. 96 *ddm1/ddm1* T2 plants from 6 independent T1 lines (16 x 6) were isolated by genotyping and phenotyping was performed, followed by confirmation of the hairpin construct insertion by PCR. Of the 96 examined plants, the number of plants which had the hairpin construct were; 71 (*hp1*), 75 (*hp2*), 76 (*hp3*), 66 (*hp4*), 74 (*hp5*), 67 (*hp6*), 72 (*hp7*), 74 (*hp8*). For phenotyping, 9-week-old plants were used for assessing height and fertility, and 7-week-old plants for the flower phenotype. 50 flowers were analyzed for each plant, and overall fertility was estimated based on seed availability (Sterile; 1-10 seeds / plant) and primary developing silique length (3-5 mm (approx. 1-5 seeds / silique); 5-7 mm (approx. 5-10 seeds / silique); 7-9 mm (approx. 10-15 seeds / silique); 9-11 mm (approx. 15-20 seeds / silique); >11 mm (> 20 seeds / silique)). Note that the *ddm1/ddm1* plants which segregated in T2 without hairpins were all sterile and did not show suppression for the height and flower phenotypes, and *hp5* suppressors were fertile at least for three generations after the plants become *ddm1*/*ddm1*, although the fertility was reduced more in later generations. Because *hp5* showed the strongest suppression, we subsequently isolated a T3 homozygous *hp5* insertion line with the heterozygous *DDM1* mutation, and used T3 *ddm1/ddm1* hp5 suppressors for RT-PCR, ChIP-seq and cytogenetics. For construction of *Cen5-ATHILA5* overexpressing plants, the *Cen5-ATHILA5* element or its ORF AT5G31927 were cloned into pMDC45 expression vector at the KpnI-SpeI site and the vectors were transformed into *rdr1;2;6* and approximately 16 T1 plants were phenotyped. The images and qRT-PCR data for the overexpressing lines were taken in selfed T2 plants.

### rdr1 rdr2 rdr6 ddm1 suppressor analysis

Seeds of *rdr1 rdr2 rdr6 DDM1/ddm1* were mutagenized with EMS (Ethyl-methanesulfonate) and *DDM1/ddm1* plants (approximately n=500) were selected by genotyping of the M1 generation. In M2, *rdr1;2;6 ddm1* plants with rescued sterility and floral defects were isolated by checking approximately 3000 M2 plants showing curly leaf and short stature phenotypes, followed by confirmation of the *ddm1* homozygous mutation by genotyping. EMS-induced SNPs in *rdr1;2;6 ddm1* suppressors were identified by whole genome sequencing (Illumina Hiseq2000). Suppressors’ parental M2 seeds (*rdr1 rdr2 rdr6 DDM1/ddm1* bearing the heterozygous suppressor mutation) were planted to segregate suppressors and non-suppressors in the same M3 progeny, allowing us to perform CAPS analysis. 15 suppressors and 45 non-suppressors were analyzed for each suppressor. SNPs in the centromeric region of chromosome 5 and restriction enzymes used for CAPS analysis are as follows: 10483242 G to A and PacI (*2-69*), 11316097 G to A and Hpy188I (*2-20*), 13349168 G to A and HhaI (*3-72*), 13818243 C to T and AflII (*3-75*).

### RNA analysis

10ug of total RNA was used for electrophoresis on 15% Acrylamide Urea-TBE gel. Separated RNA was transferred onto Hybond-NX membrane (GE Healthcare) and the membrane was crosslinked with EDC (1-ethyl-3-(3-dimethylaminopropyl)carbodiimide). RNA probes for detecting *ATHILA*-derived small RNA were generated in vitro as recommended by the manufacturer (Ambion). To prepare a probe for miR159 detection, its complementary oligo nucleotide DNA was labeled with radioactive phosphate (Perkin Elmer). For quantitative RT-PCR 1ug of total RNA was treated with 5 Units of DNase I (Takara) and cDNA synthesized with SuperScriptIII (Life Technologies) was used for the subsequent qPCR analysis.

### Whole genome bisulfite sequencing

Whole genome bisulfite sequencing was performed as described previously^18^. Reads were mapped using Bismark. Initially, hypermethylated regions common to all 4 suppressors were identified by genome browsing, but robustness was then assessed by DMR analysis. For analyzing Differentially Methylated Regions (DMRs), total DNA Methylation levels in 300bp were calculated by summing all CG/CHG/CHH methylation levels. Of the regions retaining less than 40% of DNA methylation levels in *rdr1;2;6 ddm1* compared with those in *rdr1;2;6*, the regions recovering more than 60% in suppressors were sorted as hyper-methylated DMRs in suppressors.

### Chromatin immunoprecipitation sequencing

ChIP-seq were perfomed as described previously with some modifications^55^. For the modified steps, the ground cells were further broken up by a dounce homogenizer, and Bioruptor (Diagenode) was used to shear chromatin. The antibody against H3K9me2 was obtained from Abcam (ab1220). ChIP-seq libraries were made by NEB Next Ultra II DNA Library Prep Kit. FASTQ files were quality-filtered using SICKLE, and mapping was performed using Bowtie2^56^. The files for data analysis (BEDGRAPH) were generated using samtools^57^ and bedtools^58^. Duplicate reads were kept as H3K9me2 is enriched at repetitive or multi-copy elements, but conclusions were unchanged regardless of removing duplicate reads. MACS2^59^ was used for peak calling (q value = 0.01).

### Long-read DNA sequencing (ONT) and methylation base-calling

DNA was extracted from approximately 100 mg of young plant tissue from *rdr1;2;6 ddm1, rdr1;2;6 Cen5-ATHILA5 ddm1* (*hp5*) and their corresponding DDM1 wild-type siblings with the DNeasy Plant Pro Kit (Qiagen). From each sample, 600 ng – 1 ug of purified DNA was taken as input for ligation library preparation with the Native Barcoding Kit 24 v14 (ONT – SQK-NBD114.24) with equimolar pooling following barcode ligation. 35 ng of the multiplexed library was sequenced on an R10.4.1 PromethION flow cell.

Standard and modified (5mC) base calling and alignment to the Col-CEN v1.2 reference was carried out simultaneously with bonito 0.6.2 (ONT) using the dna_r10.4.1_e8.2_400bps_sup@v3.5.2 and 5mC_all_context_sup_r1041_e82 models, producing a modbam output file. The modbam was converted to a bedMethyl file with modkit v0.1.4 (ONT) “pileup--combine-mods”. Calls at positions with >= 3 reads were retained and the remaining calls were split by cytosine context (CpG, CHG, CHH) using modkit motif-bed and bedtools v2.31.0^58^ intersect. Methylation ratios at each position were scaled to the [0-1] interval, and ratios on the (-) reference strand were multiplied by -1 before conversion to BigWig format.

### Cytogenetics

1 week-old seedlings were soaked in 1mg/ml of DAPI solution containing 0.1% Triton X-100 for 10 min at room temperature. DAPI-stained chromosomes were analyzed with ZEISS microscopy. To calculate proportion of chromocenter signals in nucleus, DAPI signals from 30 chromocenters were analyzed by Image J^38^. DNA FISH was performed as described previously^35^. For preparing probes, 2 contiguous BAC clones were used to detect each chromosome: T1F9 F11P17 (Chr 1), T2G17 F11A3 (Chr 2), MIPN9 MIMB12 (Chr 3), F6I7 F13M23 (Chr 4), MINC6 K19P17 (Chr 5, Fig. 4a) and T1G16 T1N24 (Chr 5, Fig. 6b). Probes were labeled by nick translation with Cy3-dUTP or Cy5-dUTP as recommended by the supplier (Promokine). Fluorescent signals were analyzed by confocal microscopy. Immunofluorescence experiments were performed as described previously^35^. The antibody used for detecting H3K9me2 was ab1220 (Abcam). 30 chromocenters were analyzed to measure the ratio of H3K9me2 to DAPI, and the measurement was performed with Image J.

## Acknowledgements

We thank the late Simon Chan and Taku Sasaki for sharing seeds; Soichi Inagaki, Mayumi Takahashi and Akiko Terui for experimental support; Paul Fransz for training in cytogenetics, and Joe Simorowski, Benjamin Roche and Jean-Sébastien Parent for helpful suggestions. A. S. was supported by JSPS postdoctoral fellowships. Research in the Martienssen Laboratory is supported by a grant from the National Institutes of Health (RO1GM067014) and by the Howard Hughes Medical Institute. D. G. was supported by a Marie Curie fellowship (REP-658900-2) and a grant from the Agence Nationale de la Recherche (ANR-12-BCV2-0013). T. K. was supported by the Japanese Ministry of Education, Culture, Sports, Science, and Technology (26221105 and 15H05963).

## Author Contributions

R.A.M., A.S., I.H. and T.K. designed the study. A.S., E.E., J.L. and D.G. performed experiments. A.S., J.C., E.E. and D.G. analyzed data. R.A.M. and A.S. prepared the manuscript.

## Data Availability

Sequence data that support the findings of this study have been deposited in Gene Expression Omnibus with the accession codes GSE132005 (https://www.ncbi.nlm.nih.gov/geo/query/acc.cgi?acc=GSE132005).

## References

1. Steiner, F. A. & Henikoff, S. Diversity in the organization of centromeric chromatin. Curr. Opin. Genet. Dev. 31, 28–35 (2015).

2. Presting, G. G. Centromeric retrotransposons and centromere function. Curr. Opin. Genet. Dev. 49, 79–84 (2018).

3. Birchler, J. A. & Han, F. Barbara McClintock’s Unsolved Chromosomal Mysteries: Parallels to Common Rearrangements and Karyotype Evolution. Plant Cell 30, 771– 779 (2018).

4. Matzke, M. A. & Mosher, R. A. RNA-directed DNA methylation: an epigenetic pathway of increasing complexity. Nat. Rev. Genet. 15, 394–408 (2014).

5. Zhou, M. & Law, J. A. RNA Pol IV and V in gene silencing: Rebel polymerases evolving away from Pol II’s rules. Curr. Opin. Plant Biol. 27, 154–164 (2015).

6. Pal-Bhadra, M. et al. Heterochromatic silencing and HP1 localization in Drosophila are dependent on the RNAi machinery. Science 303, 669–672 (2004).

7. Gu, S. G. et al. Amplification of siRNA in Caenorhabditis elegans generates a transgenerational sequence-targeted histone H3 lysine 9 methylation footprint. Nat. Genet. 44, 157–164 (2012).

8. Volpe, T. A. et al. Regulation of heterochromatic silencing and histone H3 lysine-9 methylation by RNAi. Science 297, 1833–1837 (2002).

9. Hall, I. M., Noma, K.-I. & Grewal, S. I. S. RNA interference machinery regulates chromosome dynamics during mitosis and meiosis in fission yeast. Proc. Natl. Acad. Sci. U. S. A. 100, 193–198 (2003).

10. Volpe, T. et al. RNA interference is required for normal centromere function in fission yeast. Chromosome Res. 11, 137–146 (2003).

11. May, B. P., Lippman, Z. B., Fang, Y., Spector, D. L. & Martienssen, R. A. Differential regulation of strand-specific transcripts from Arabidopsis centromeric satellite repeats. PLoS Genet. 1, e79 (2005).

12. Vongs, A., Kakutani, T., Martienssen, R. A. & Richards, E. J. Arabidopsis thaliana DNA methylation mutants. Science 260, 1926–1928 (1993).

13. Borges, F. & Martienssen, R. A. The expanding world of small RNAs in plants. Nat. Rev. Mol. Cell Biol. 16, 727–741 (2015).

14. Slotkin, R. K. et al. Epigenetic reprogramming and small RNA silencing of transposable elements in pollen. Cell 136, 461–472 (2009).

15. Lippman, Z. et al. Role of transposable elements in heterochromatin and epigenetic control. Nature 430, 471–476 (2004).

16. Fu, F.-F., Dawe, R. K. & Gent, J. I. Loss of RNA-directed DNA Methylation in Maize Chromomethylase and DDM1-type Nucleosome Remodeler Mutants. Plant Cell 30, tpc.00053.2018 (2018).

17. McCue, A. D., Nuthikattu, S. & Slotkin, R. K. Genome-wide identification of genes regulated in trans by transposable element small interfering RNAs. RNA Biol. 10, 1379–1395 (2013).

18. Creasey, K. M. et al. miRNAs trigger widespread epigenetically activated siRNAs from transposons in Arabidopsis. Nature 508, 411–415 (2014).

19. Wang, X.-B. et al. RNAi-mediated viral immunity requires amplification of virus-derived siRNAs in *Arabidopsis thaliana*. Proc. Natl. Acad. Sci. U. S. A. 107, 484–489 (2010).

20. Zemach, A. et al. The arabidopsis nucleosome remodeler DDM1 allows DNA methyltransferases to access H1-containing heterochromatin. Cell 153, 193–205 (2013).

21. Teixeira, F. K. et al. A role for RNAi in the selective correction of DNA methylation defects. Science 323, 1600–1604 (2009).

22. Sasaki, T., Kobayashi, A., Saze, H. & Kakutani, T. RNAi-independent de novo DNA methylation revealed in Arabidopsis mutants of chromatin remodeling gene DDM1. Plant J. 70, 750–758 (2012).

23. Saze, H. & Kakutani, T. Heritable epigenetic mutation of a transposon-flanked Arabidopsis gene due to lack of the chromatin-remodeling factor DDM1. EMBO J. 26, 3641–3652 (2007).

24. Henderson, I. R. & Jacobsen, S. E. Tandem repeats upstream of the Arabidopsis endogene SDC recruit non-CG DNA methylation and initiate siRNA spreading. Genes and Development 22, 1597–1606 (2008).

25. Kankel, M. W. et al. Arabidopsis MET1 cytosine methyltransferase mutants. Genetics 163, 1109–1122 (2003).

26. Lippman, Z., May, B., Yordan, C., Singer, T. & Martienssen, R. A. Distinct mechanisms determine transposon inheritance and methylation via small interfering RNA and histone modification. PLoS Biol. 1, (2003).

27. Copenhaver, G. P. et al. Genetic definition and sequence analysis of Arabidopsis centromeres. Science 286, 2468–2474 (1999).

28. Jacobsen, S. E. & Meyerowitz, E. M. Hypermethylated SUPERMAN epigenetic alleles in arabidopsis. Science 277, 1100–1103 (1997).

29. Soppe, W. J. J. et al. The late flowering phenotype of fwa mutants is caused by gain-of-function epigenetic alleles of a homeodomain gene. Mol. Cell 6, 791–802 (2000).

30. Finnegan, E. J., Peacock, W. J. & Dennis, E. S. DNA methylation, a key regulator of plant development and other processes. Curr. Opin. Genet. Dev. 10, 217–223 (2000).

31. Ravi, M. et al. The rapidly evolving centromere-specific histone has stringent functional requirements in Arabidopsis thaliana. Genetics 186, 461–471 (2010).

32. Naish, M. et al. The genetic and epigenetic landscape of the Arabidopsis centromeres. Science 374, (2021).

33. Wlodzimierz, P. et al. Cycles of satellite and transposon evolution in Arabidopsis centromeres. Nature 618, 557–565 (2023).

34. Lee, S. C. et al. Arabidopsis retrotransposon virus-like particles and their regulation by epigenetically activated small RNA. Genome Res. 30, 576–588 (2020).

35. Yelagandula, R. et al. The histone variant H2A.W defines heterochromatin and promotes chromatin condensation in Arabidopsis. Cell 158, 98–109 (2014).

36. Mathieu, O., Reinders, J., Čaikovski, M., Smathajitt, C. & Paszkowski, J. Transgenerational Stability of the Arabidopsis Epigenome Is Coordinated by CG Methylation. Cell 130, 851–862 (2007).

37. Carmena, M., Wheelock, M., Funabiki, H. & Earnshaw, W. C. The chromosomal passenger complex (CPC): from easy rider to the godfather of mitosis. Nat. Rev. Mol. Cell Biol. 13, 789–803 (2012).

38. Batzenschlager, M. et al. Arabidopsis MZT1 homologs GIP1 and GIP2 are essential for centromere architecture. Proc. Natl. Acad. Sci. U. S. A. 112, 8656–8660 (2015).

39. Smith, S. J., Osman, K. & Franklin, F. C. H. The condensin complexes play distinct roles to ensure normal chromosome morphogenesis during meiotic division in Arabidopsis. Plant J. 80, 255–268 (2014).

40. Simon, L., Voisin, M., Tatout, C. & Probst, A. V. Structure and Function of Centromeric and Pericentromeric Heterochromatin in Arabidopsis thaliana. Front. Plant Sci. 6, 1049 (2015).

41. Shibata, F. & Murata, M. Differential localization of the centromere-specific proteins in the major centromeric satellite of Arabidopsis thaliana. J. Cell Sci. 117, 2963–2970 (2004).

42. Demidov, D. et al. Altered expression of Aurora kinases in Arabidopsis results in aneu- and polyploidization. Plant J. 80, 449–461 (2014).

43. Ebbs, M. L. & Bender, J. Locus-specific control of DNA methylation by the Arabidopsis SUVH5 histone methyltransferase. Plant Cell 18, 1166–1176 (2006).

44. Yelina, N. E. et al. DNA methylation epigenetically silences crossover hot spots and controls chromosomal domains of meiotic recombination in Arabidopsis. Genes and Development 29, 2183–2202 (2015).

45. Gutbrod, M. J. et al. Dicer promotes genome stability via the bromodomain transcriptional co-activator BRD4. Nat. Commun. 13, 1001 (2022).

46. Unoki, M., Funabiki, H., Velasco, G., Francastel, C. & Sasaki, H. CDCA7 and HELLS mutations undermine nonhomologous end joining in centromeric instability syndrome. J. Clin. Invest. 129, 78–92 (2019).

47. Marasco, L. E. et al. Counteracting chromatin effects of a splicing-correcting antisense oligonucleotide improves its therapeutic efficacy in spinal muscular atrophy. Cell 185, 2057–2070.e15 (2022).

48. Karpen, G. H. & Allshire, R. C. The case for epigenetic effects on centromere identity and function. Trends Genet. 13, 489–496 (1997).

49. Koornneef, M. & Van der Veen, J. H. Trisomics inArabidopsis thaliana and the location of linkage groups. Genetica 61, 41–46 (1983).

50. Zhang, W., Lee, H.-R., Koo, D.-H. & Jiang, J. Epigenetic modification of centromeric chromatin: hypomethylation of DNA sequences in the CENH3-associated chromatin in Arabidopsis thaliana and maize. Plant Cell 20, 25–34 (2008).

51. Wells, J. N. & Feschotte, C. A Field Guide to Eukaryotic Transposable Elements. Annu. Rev. Genet. 54, 539–561 (2020).

52. Gent, J. I., Wang, N. & Dawe, R. K. Stable centromere positioning in diverse sequence contexts of complex and satellite centromeres of maize and wild relatives. Genome Biol. 18, 121 (2017).

53. Niki, H. Schizosaccharomyces japonicus: the fission yeast is a fusion of yeast and hyphae. Yeast 31, 83–90 (2014).

54. Tanabe, S. et al. A novel cytochrome P450 is implicated in brassinosteroid biosynthesis via the characterization of a rice dwarf mutant, dwarf11, with reduced seed length. Plant Cell 17, 776–790 (2005).

55. Inagaki, S. et al. Gene-body chromatin modification dynamics mediate epigenome differentiation in Arabidopsis. EMBO J. 36, 970–980 (2017).

56. Langmead, B. & Salzberg, S. L. Fast gapped-read alignment with Bowtie 2. Nat. Methods 9, 357–359 (2012).

57. Li, H. et al. The Sequence Alignment/Map format and SAMtools. Bioinformatics 25, 2078–2079 (2009).

58. Quinlan, A. R. & Hall, I. M. BEDTools: A flexible suite of utilities for comparing genomic features. Bioinformatics 26, 841–842 (2010).

59. Zhang, Y. et al. Model-based analysis of ChIP-Seq (MACS). Genome Biol. 9, (2008).

60. Schneider, C. A., Rasband, W. S. & Eliceiri, K. W. NIH Image to ImageJ: 25 years of image analysis. Nat. Methods 9, 671–675 (2012).

