## Supplementary Figures for "Retrotransposon addiction promotes centromere function via epigenetically activated small RNAs"

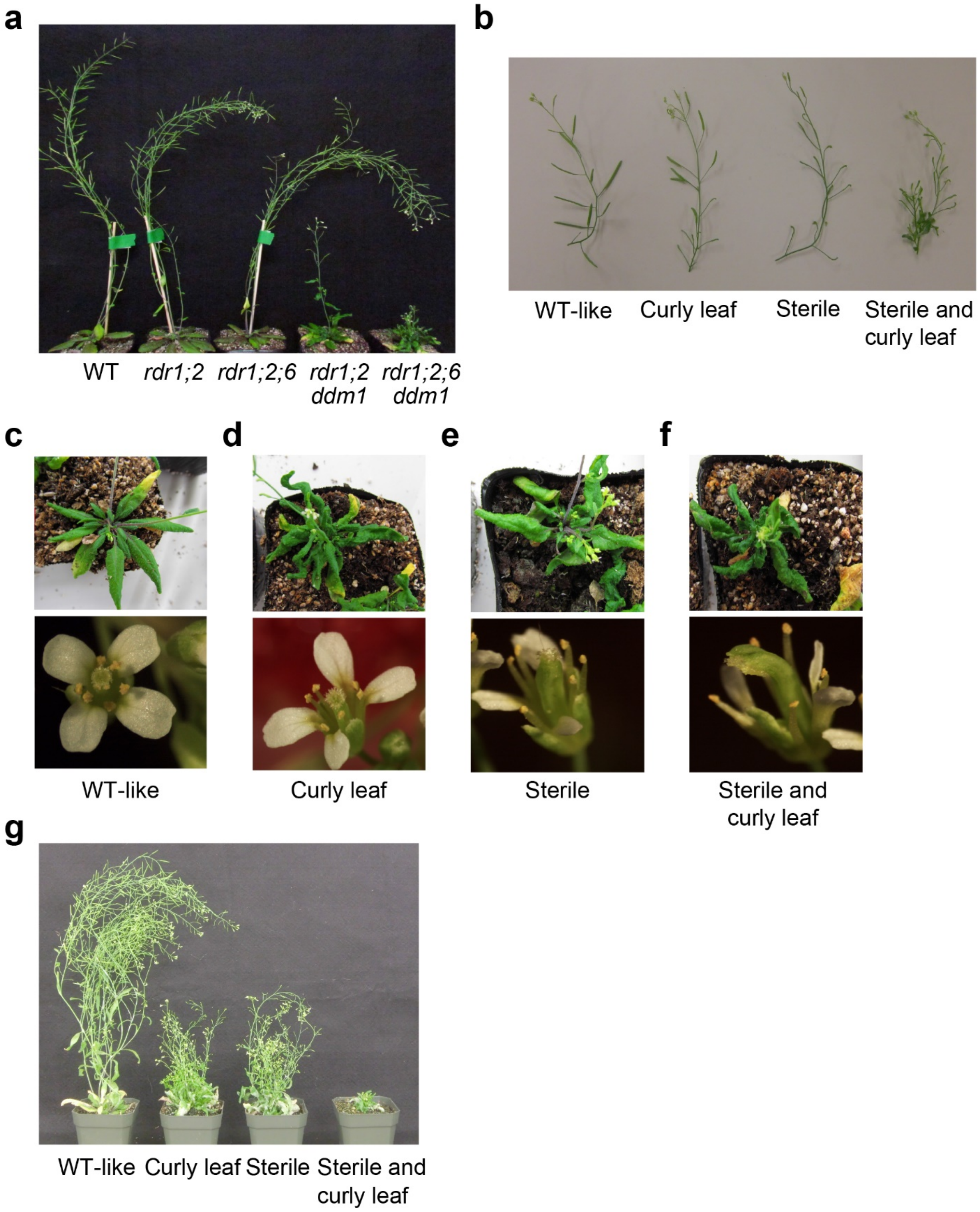


**Extended Data Figure 1. Phenotypes of *rdr1;2;6 ddm1* and of *ddm1* epiRILs in an *rdr1;2;6* background.**

**a**, Plant stature phenotypes in the indicated genotypes. **b-g**, The phenotypes of *ddm1* epiRILs were classified into 4 groups (WT-like, Curly leaf, Sterile, Sterile and Curly leaf). Panels show (**b**) siliques, (**c-f**) flowers and leaves, and (**g**) stature of each group.

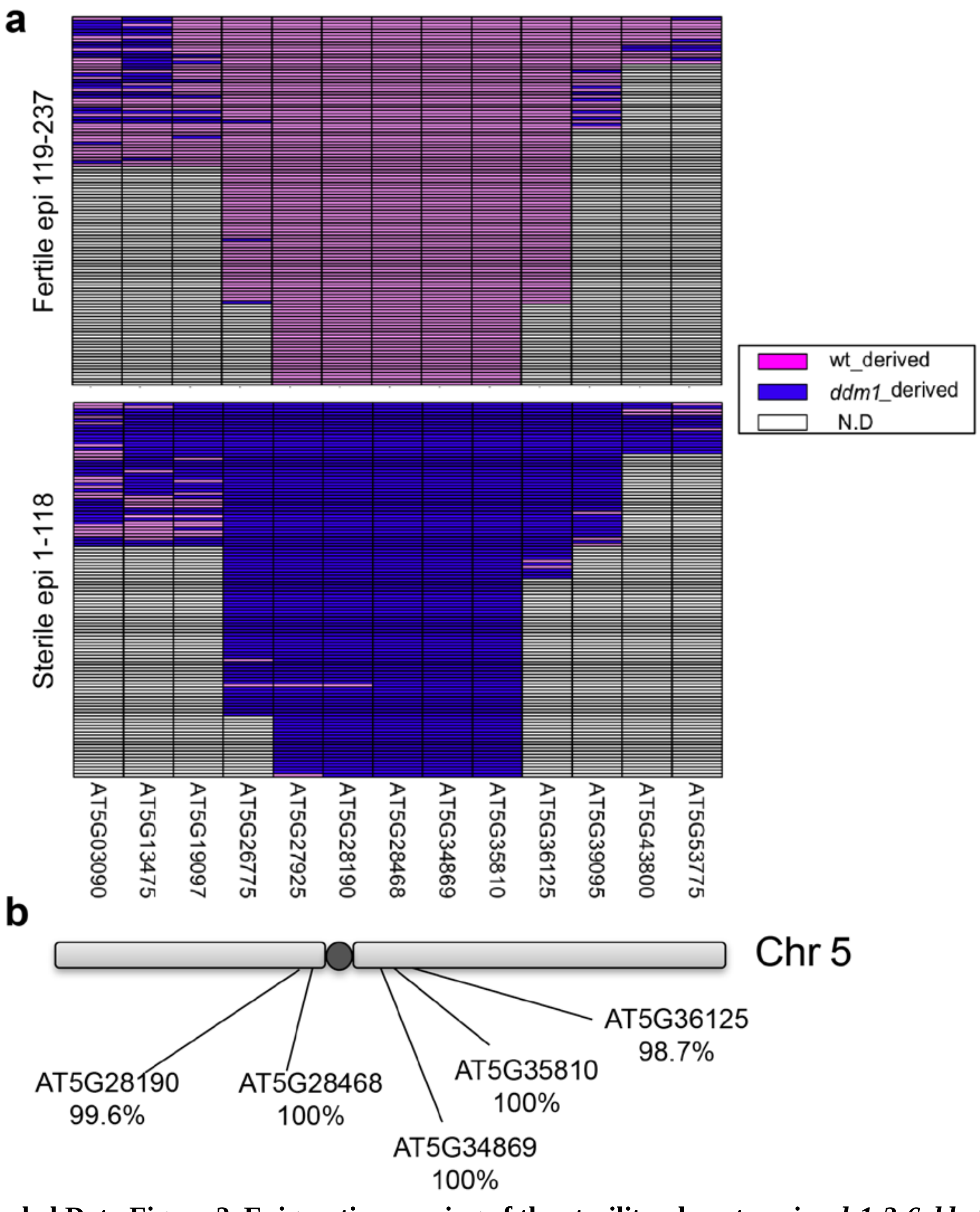


**Extended Data Figure 2. Epigenetic mapping of the sterility phenotype in *rdr1;2;6 ddm1*.**
**a**, DNA was extracted from fertile and sterile *ddm1* epigenetic recombinant lines (sterile;epi 1-118, fertile;epi 119-237) in *rdr1;2;6* background, and DNA methylation at the indicated transposable elements (AT5G03090-AT5G53775) was assessed by McrBC-based PCR analysis (see methods). Upper and lower panels indicate methylation maps of fertile and sterile epigenetic recombinant lines, respectively. Chromosomal regions derived from WT and *ddm1* are colored in pink and blue, respectively. **b**, An epigenetic linkage map of the sterility phenotype in *rdr1;2;6*. Linkage between DNA hypomethylation with the sterile phenotype is indicated below TE gene names.

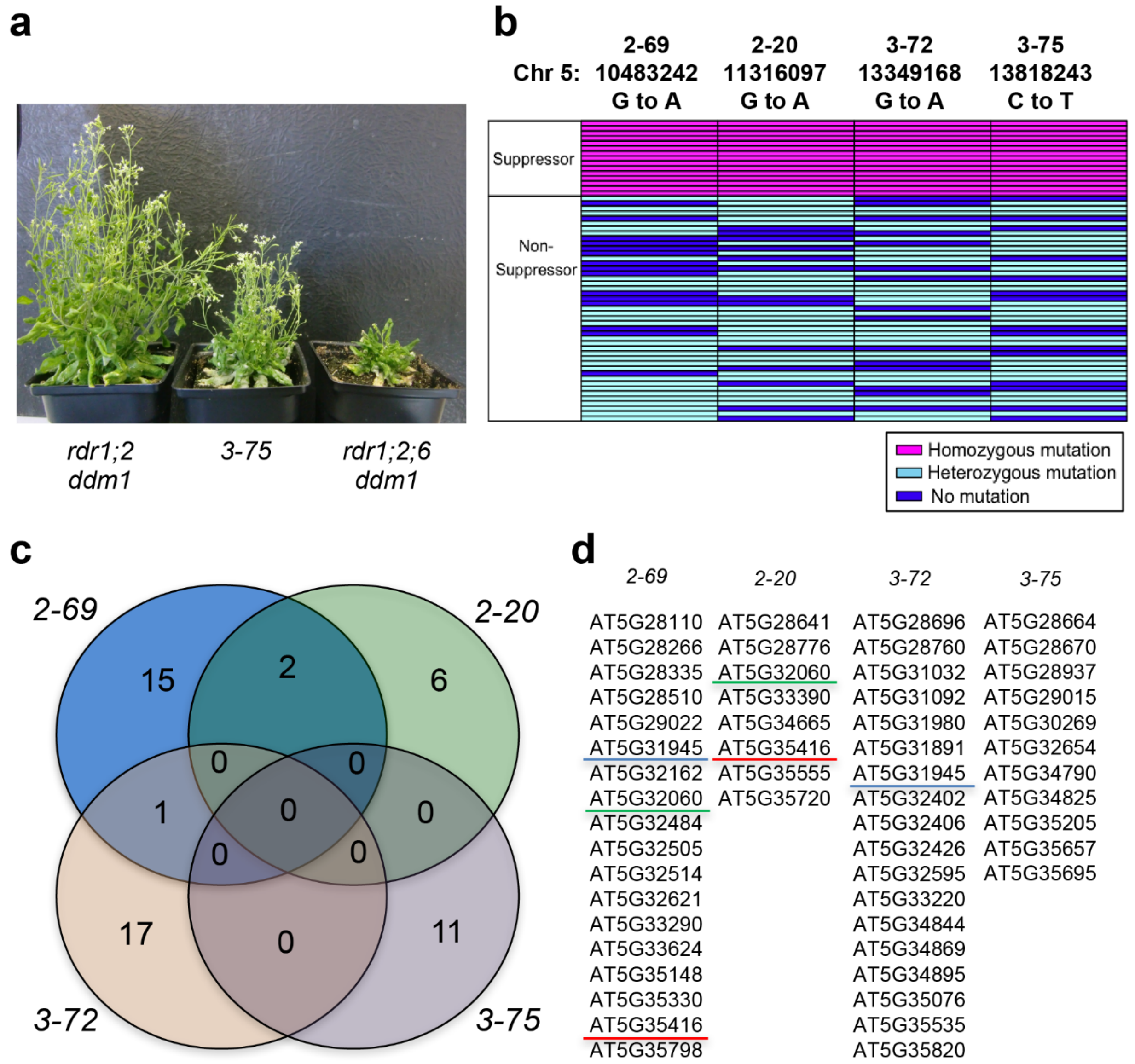


**Extended Data Figure 3. Genetic mapping of linked mutations in EMS suppressor lines.**
**a**, EMS suppressor 3-75 rescued the phenotype of *rdr1;2;6 ddm1*. **b**, CAPS analysis using EMS-induced SNPs was performed in M2 progeny segregating suppressors and non-suppressors. We focused on chromosome 5 centromeric region where the sterility defect mapped (Extended Data Figure 2). SNPs used in the analysis are shown above the panel. Cells in the table are colored by pink, light blue and blue, to indicate individuals bearing homozygous SNP, heterozygous SNP and no SNP, respectively. Each suppressor was recessive and tightly linked to Cen5. **c**, Venn diagram of mutations detected on chromosome 5 centromeric regions in EMS suppressors. **d**, A list of mutations introduced into chromosome 5 centromeric regions in *rdr1;2;6 ddm1* suppressors. Genes underlined with the same color represent mutated genes in more than one suppressor.

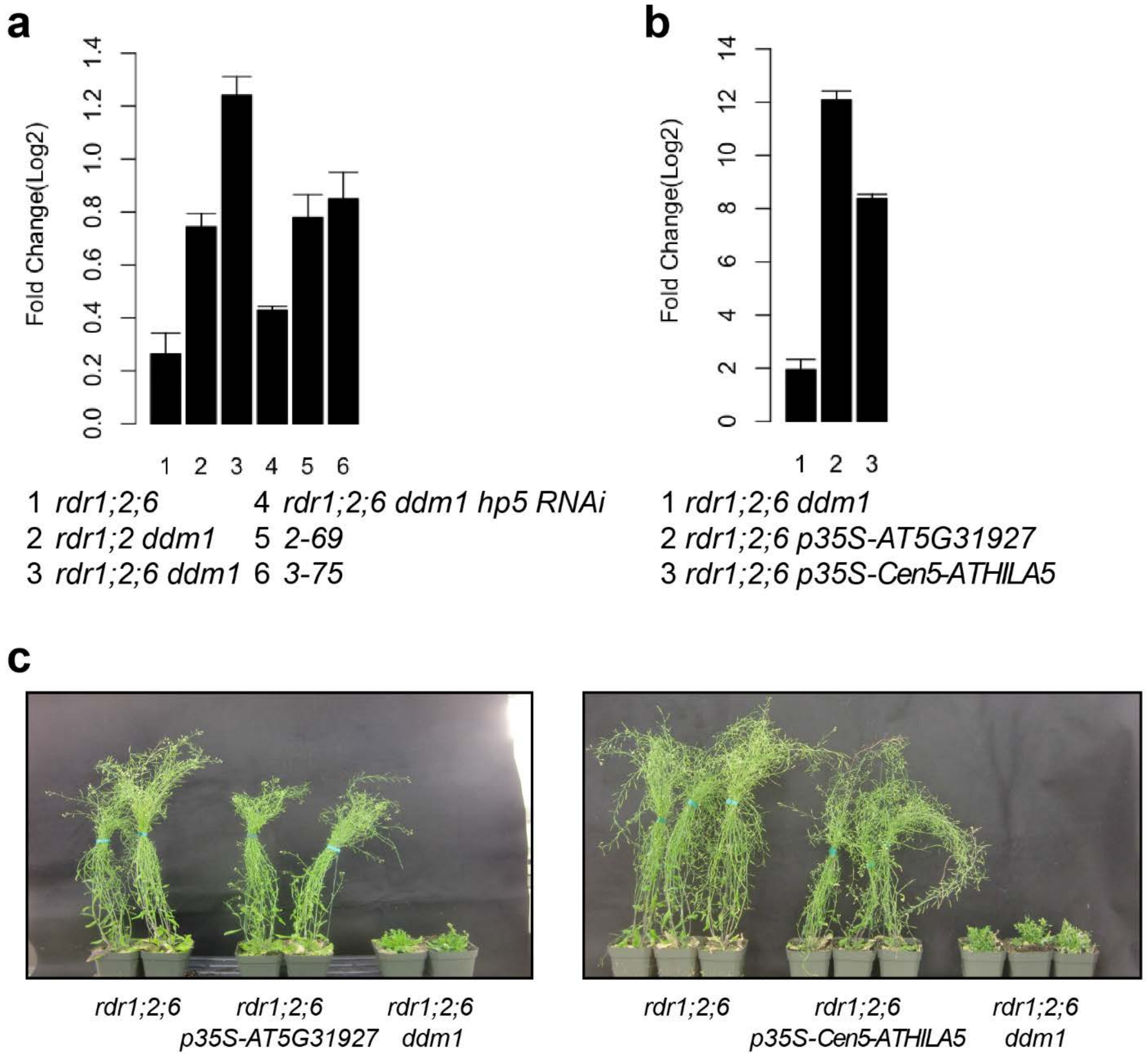


**Extended Data Figure 4. Overexpression of *Cen5-ATHILA5* does not cause developmental phenotypes in *rdr1;2;6* triple mutants.**

**a**, Expression levels of *Cen5-ATHILA5* in the indicated plants were analyzed by RT-qPCR. Signals were normalized with *Cen5-ATHILA5* in WT. Bars represent standard error. **b**, RT-qPCR analysis for *Cen5-ATHILA5* in the plants overexpressing *AT5G31927* and *Cen5-ATHILA5*. Signals were normalized with *Cen5-ATHILA5* in WT. **c**, Photos of 6-week-old *rdr1;2;6* plants overexpressing *AT5G31927* (left panel) and overexpressing *Cen5-ATHILA5* (right panel).

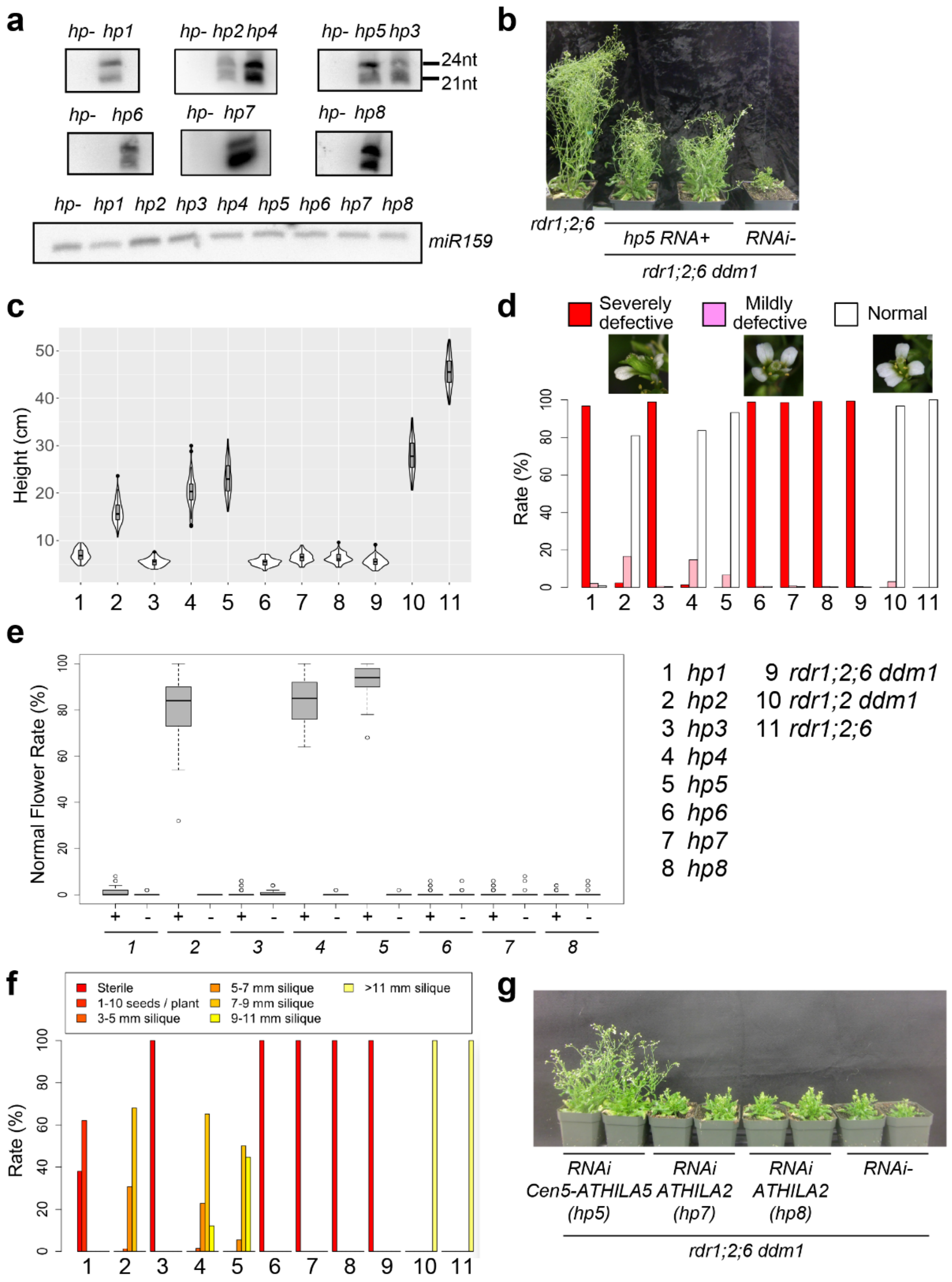


**Extended Data Figure 5. Hairpin suppressors of fertility and stature defects in *rdr1;2;6 ddm1***
**a**, *ATHILA5* and *ATHILA2* small RNAs were detected by Northern blot in the absence (*hp-*) or presence of hairpins shown in Fig. 3 (*hp1-6*: *Cen5-ATHILA5*; *hp7-8*: *ATHILA2*). As a loading control, abundantly expressed miR159 was detected on a different gel. **b**, Phenotypic suppression by *hp5* in *rdr1;2;6 ddm1*. **c-f**, The effects of *Cen5-ATHILA5* and *ATHILA2* hairpins on the *rdr1;2;6 ddm1* phenotypes: (**c**) height; (**d, e**) normal flowers and (**f**) fertility (silique length). The labeled numbers below the figures indicate the mutants shown next to (**e**). **g**, 6-week-old *rdr1;2;6 ddm1* plants expressing *Cen5-ATHILA5* (*hp5*) or *ATHILA2* hairpins (*hp7* and *hp8*).

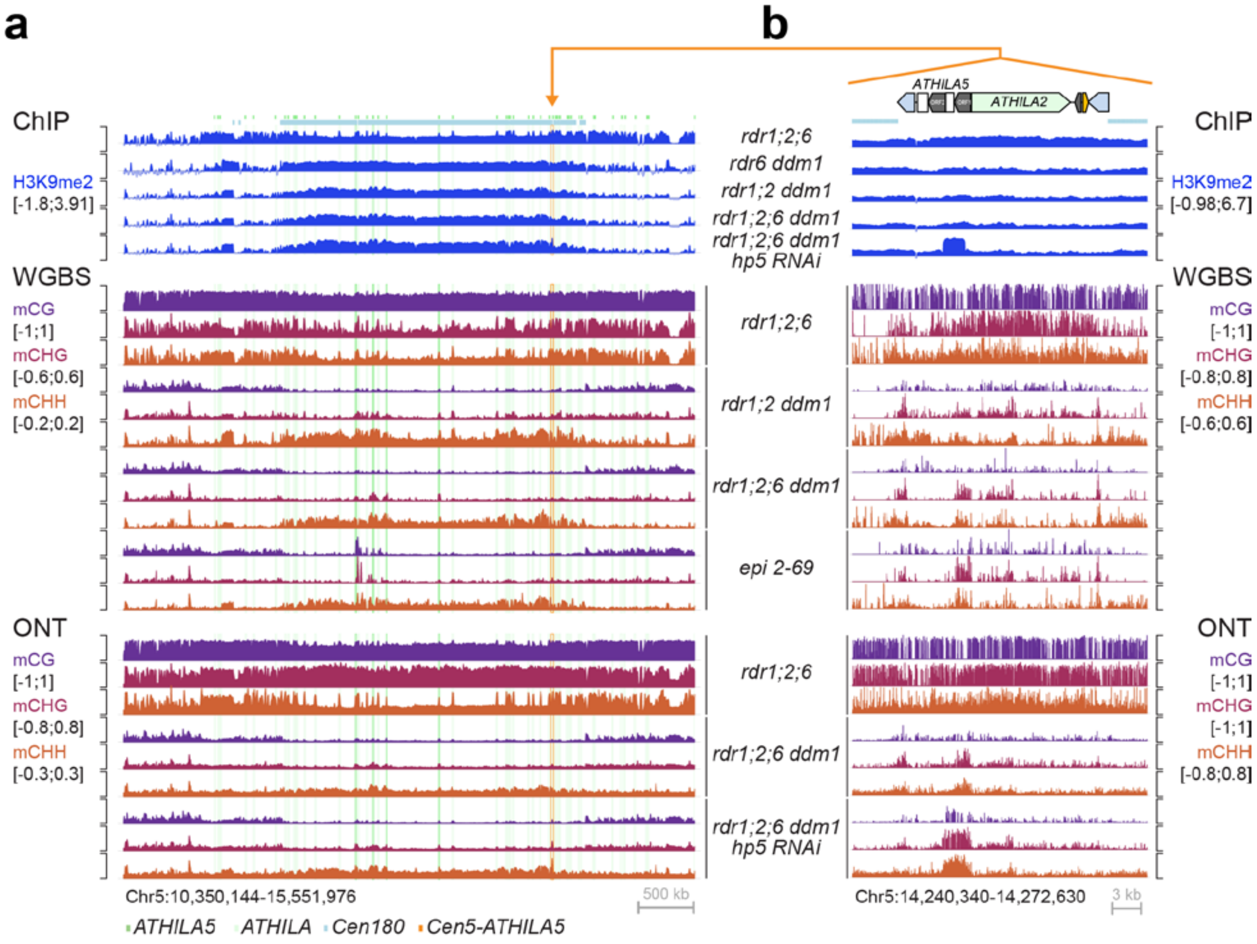


**Extended Data Figure 6. Comparison of DNA methylation and H3K9me2 levels at Chromosome 5 pericentromere in the different genotypes.**
**a, b**, Browser screenshots of (**a**) the centromeric region of Chromosome 5 and (**b**) *Cen5-ATHILA5*, showing H3K9me2 tracks (log2[IP/Input]), DNA methylation (mC/C in each sequence context) from whole-genome bisulfite sequencing (WGBS), and DNA methylation called with modbase2 from long reads Oxford Nanopore Technologies (ONT). The loss of H3K9me2 and DNA methylation in the *rdr1;2;6 ddm1* mutant is partially recovered at the *Cen5-ATHILA5* with expression of the RNAi hairpin *hp5*. Values are averaged in windows of (**a**) 5 kb or (**b**) 10 bp.

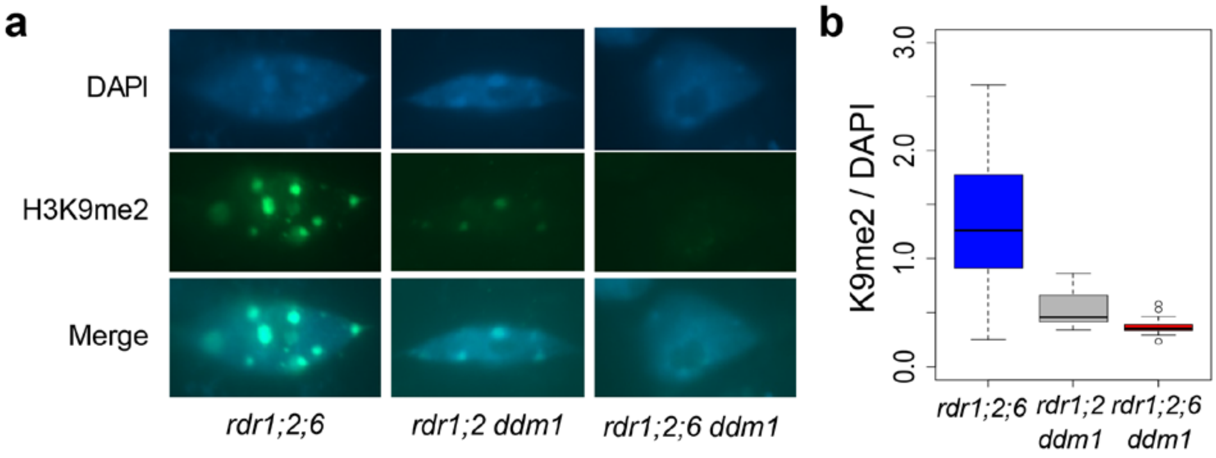


**Extended Data Figure 7. Reduced H3K9me2 at chromocenters in *rdr1 rdr2 rdr6 ddm1*.**
**a**, Immunofluorescence of H3K9me2 in mature leaf nuclei was observed in the indicated genotypes, and nuclei were counterstained with DAPI (top panels). **b**, Ratios of H3K9me2 to DAPI in chromocenters of each genotype.

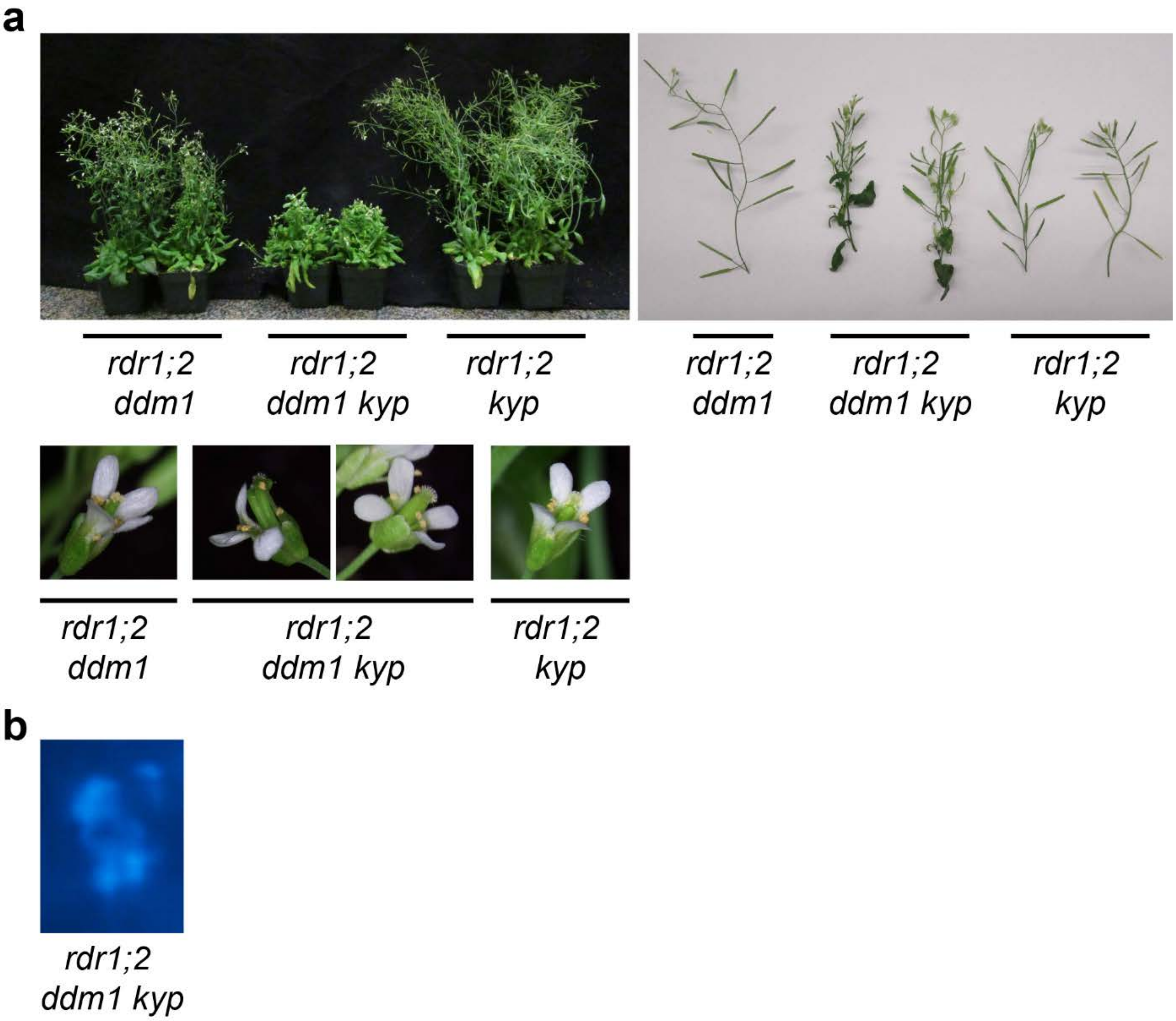


**Extended Data Figure 8. Developmental phenotypes and chromosome mis-segregation defects in *rdr1;2 ddm1 kyp*.**

**a**, Aggravated phenotypes of *rdr1;2 ddm1 kyp* compared to *rdr1;2 ddm1 and rdr1;2 kyp*. *rdr1;2 ddm1 kyp* plants exhibited partially deformed flowers (abnormal; 71%, normal; 29%) and reduced fertility. **b**, Mis-segregating chromosomes during anaphase in *rdr1;2 ddm1 kyp*.

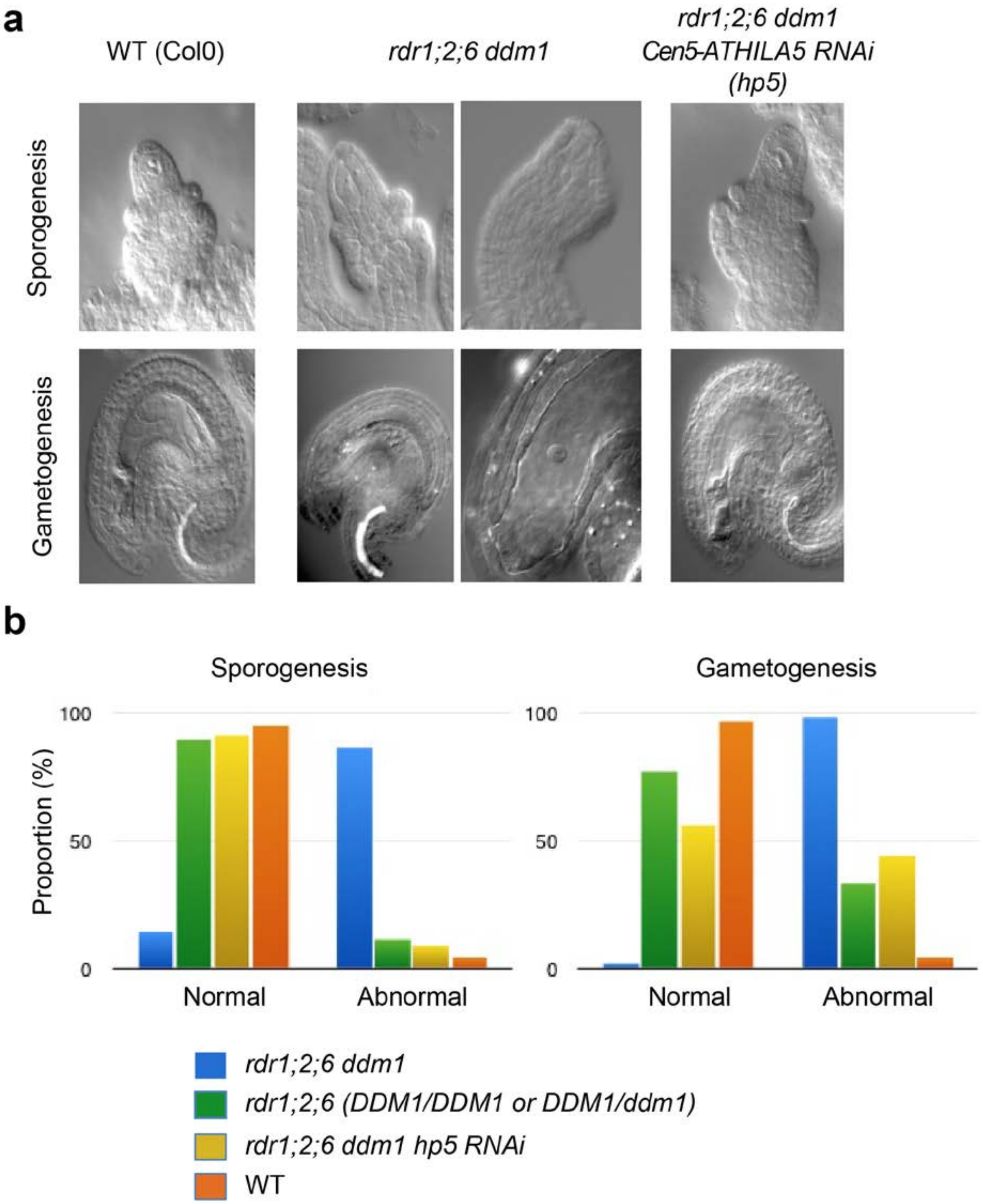


**Extended Data Figure 9. Defective female sporogenesis and gametogenesis in *rdr1;2;6 ddm1*.** **a**, Female sporogenesis and gametogenesis were analyzed by whole-mount ovule clearing in the indicated strains. In the quadruple mutant, presence of multiple megaspore mother cells (mmc) was noted in 32% of ovules during sporogenesis. Lack of a clear mmc was observed in 12% of ovules scored. Conspicuous absence of a gametophyte, or incomplete gametophytes were found in most (>76% of ovules), a phenotype which was partially rescued in the suppressor lines. **b**, Quantification of the phenotypes in the indicated strains.
